# Pathways of pathogenicity: The transcriptional stages of germination in the fatal fungal pathogen *Rhizopus delemar*

**DOI:** 10.1101/330969

**Authors:** Poppy C. S. Sephton Clark, Jose F. Muñoz, Elizabeth R. Ballou, Christina A. Cuomo, Kerstin Voelz

## Abstract

*Rhizopus delemar* is an invasive fungal pathogen, responsible for the frequently fatal disease mucormycosis. Germination, a crucial mechanism by which spores of *Rhizopus delemar* infect and cause disease, is a key developmental process that transforms the dormant spore state into a vegetative one. Understanding the molecular mechanisms which underpin this transformation may be key to controlling mucormycosis, however the regulation of germination remains poorly understood. This study describes the phenotypic and transcriptional changes which take place over the course of germination. This process is characterised by four distinct stages: dormancy, isotropic swelling, germ tube emergence and hyphal growth. Dormant spores are shown to be transcriptionally unique, expressing a subset of transcripts absent in later developmental stages. A large shift in the expression profile is prompted by the initiation of germination, with genes involved in respiration, chitin, cytoskeleton and actin regulation appearing to be important for this transition. A period of transcriptional consistency can be seen throughout isotropic swelling, before the transcriptional landscape shifts again at the onset of hyphal growth. This study provides a greater understanding of the regulation of germination and highlights processes involved in transforming *Rhizopus delemar* from a single to a multicellular organism.

**Importance:** Germination is key to the growth of many organisms, including fungal spores. Mucormycete spores exist abundantly within the environment and germinate to form hyphae. These spores are capable of infecting immunocompromised individuals, causing the disease mucormycosis Germination from spore to hyphae within patients leads to angioinvasion, tissue necrosis and often fatal infections. This study advances our understanding of how spore germination occurs in the mucormycetes, identifying processes we may be able to inhibit to help prevent or treat mucormycosis.

## Introduction

Fungal spores are found ubiquitously within the environment and are key to the dispersal and survival of many fungal species(1, 2). Spores can endure severe temperatures, desiccation, and high levels of radiation and radical exposure, conditions fatal to many other lifeforms(3). The ability to survive in harsh environments has enabled the spread of fungal spores by wind, water and animal dispersal across the globe. Once distributed, spores may stay dormant for thousands of years(4), before germination is initiated under favourable conditions.

Germination cues can include, but are not limited to, the introduction of nutrients, the presence of light, temperature modulation, changes in osmolarity, pH shifts, the removal of dormancy factors and the introduction of extracellular signalling molecules(5–15). Once germination is initiated, spores begin to swell and take up water. At a critical point, the cell polarizes(16) and hyphae emerge from the swollen spore bodies. Given the correct conditions, the transition from dormancy to vegetative hyphal growth can occur in as little as 6 hours, allowing the fungi to rapidly colonize favourable environments. Fungal spores are the infectious agents of many fungal diseases(17–19) (e.g. mucormycosis, aspergillosis, blastomycosis, cryptococcosis, coccidioidomycosis, histoplasmosis). The transition from dormancy to vegetative growth allows for the onset of disease within a host, yet we currently have a limited understanding of the molecular pathways regulating this fundamental developmental process in human pathogenic fungi(20–28).

Mucormycosis is an emerging fungal infectious disease with extremely high mortality rates, at over 90% in disseminated cases(29). Current antifungal treatments are ineffective, resulting in the reliance upon surgical debridement of infected tissues(30), often leading to long-term disability. Disease can be caused by several species of the Mucorales order, however *Rhizopus delemar*, previously known as *Rhizopus oryzae,* accounts for 70% of cases(31). Spores are the infectious agents of mucormycosis. While immunocompetent individuals control spore germination through phagocytic uptake, mucormycete spores can survive within immune effector cells, causing latent infection(32). In immunocompromised patients, inhibition of spore germination by phagocytes fails, enabling fungal growth(33). Hyphal extension within tissue leads to angioinvasion, thrombosis, tissue necrosis and eventually death(30, 34). Given the significance of spore germination in mucormycosis pathogenesis, medical interventions that target and inhibit this developmental process might improve patient prognosis. Therefore, we aimed to comprehensively characterise the transcriptional and phenotypic changes that occur over time during this process.

Phenotypic and transcriptional approaches were taken to follow the germination of *Rhizopus delemar* over time. With the previously annotated genome of *Rhizopus delemar*(35), shown to have undergone a whole genome duplication, our RNA-Seq data was analysed and used to create an updated gene set. Our data reveals a clear progression of transcriptional regulation over time, linked to observed phenotypic changes. Together, this work represents the most comprehensive analysis of the transcriptional landscape during germination in a human fungal pathogen to date.

## Results

### Phenotypic Characteristics of Germinating R. delemar

Germination is characterised by three distinct transitions: dormancy to swelling, swelling to germ tube emergence, and the switch to sustained filamentous growth. This process is common to many filamentous fungi, though the timing of germination varies between species(36). We therefore characterized the phenotypic progression of *Rhizopus delemar* strain RA 99-880 through germination by live cell imaging (Figure 1). The switch from dormancy to swelling was triggered by exposure to rich media. Swelling, characterised by an isotropic increase in size, continued for 4-6 hours (Figure 1A). Between resting and fully swollen, the average spore diameter increased from 5 μm to 13 μm (Figure 1B). Once fully swollen, germ tubes emerged from the spore bodies. Most spore bodies (75.5%, Figure 1C) produced hyphae which exceeded the diameter of the spore body in length by 4 - 5 hours. At this time point, the spores were considered fully germinated. Hyphal growth continued from 6 – 24 hours, demonstrated by increase in optical density (Figure 1D), with the average width of hyphae being 5 μm ± 1.03 μm, and the average length being 135 μm ± 30 μm.

**Figure 1:**
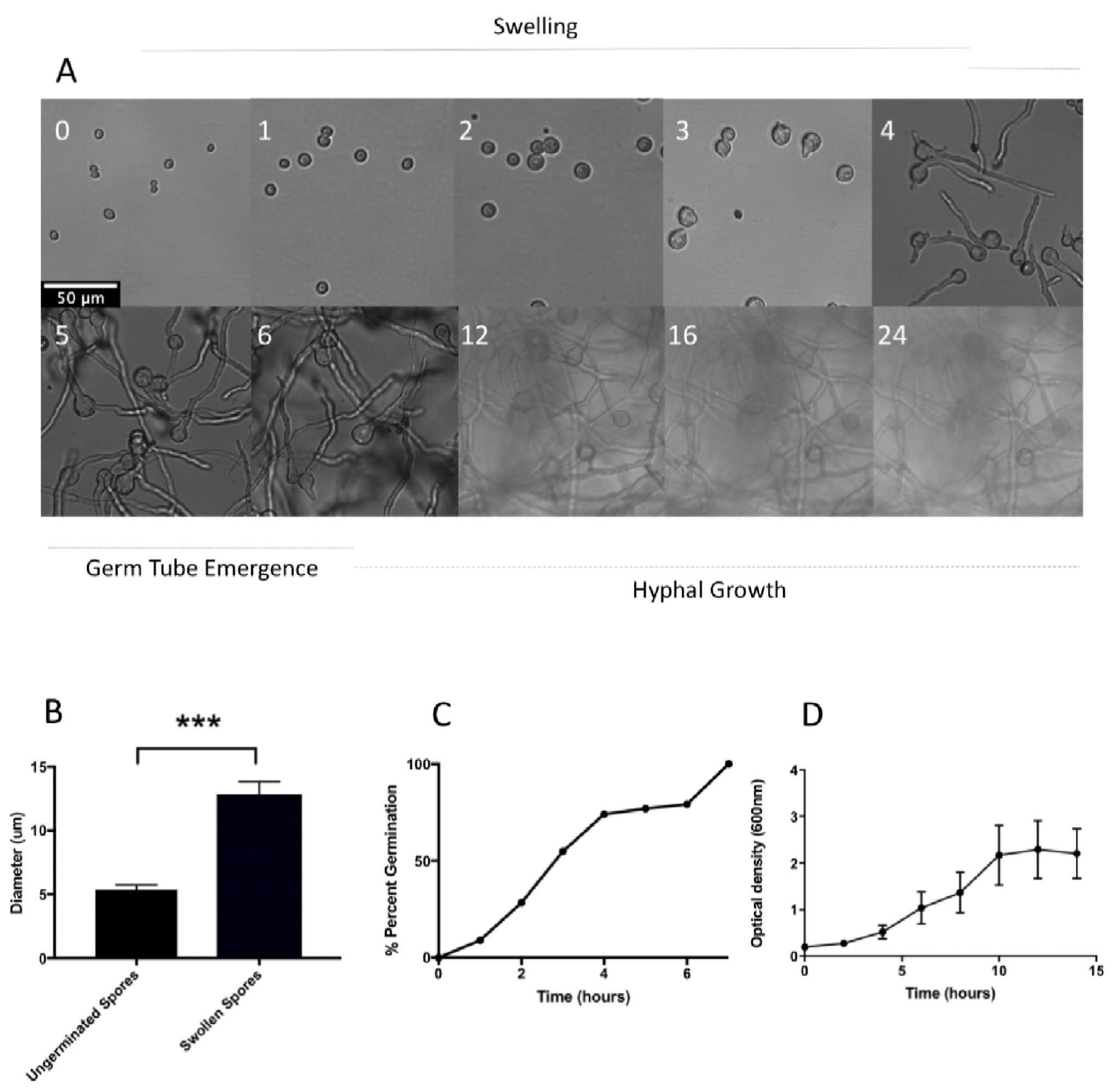
Phenotypic characterisation of germinating spores. A) Spores germinated in SAB were imaged at hours, indicated in white, post germination. Scale bar = 50 µm for all images. Micrographs representative of >3 replicate experiments are shown. B) Diameter of ungerminated spore bodies (n=3, t=0) compared to spore body size measured immediately prior to germ tube emergence for each individual spore (n=3, t=4-6). C) Spores germination as a percentage over time, determined by live cell imaging (n=3). D) Fungal mass over time, determined by optical density at 600 nm.

### Transcription Over Time: Experimental Design

Our phenotypic analysis of spore germination established the temporal pattern for the development of spores from dormancy to filamentous growth. These dramatic morphological changes require vast cellular reprogramming. In this study, we performed transcriptional analysis of each stage outlined in this process. For high-resolution capture of the transcriptional regulation of spore germination, we isolated and sequenced mRNA from resting spores (0 hours), swelling spores (1, 2, 3, 4 and 5 hours) and during filamentous growth (6, 12, 16 and 24 hours). Three biological replicates were produced for each time point, mRNA from each sample was sequenced with Illumina HiSeq technology, with 100bp paired end reads. Reads were aligned to the *R. delemar* genome(35), giving an average alignment rate of over 95% per sample, with an average of 68% (12,170 genes) of all genes expressed over all time points. We utilised our RNA-Seq data to revise the current annotation of the available *R. delemar* genome, using BRAKER 2.1.0(37) to improve gene structures and incorporate these into an updated annotation. Compared to the previous annotation(35), this updated set included 475 new predicted genes, 370 new protein family domains (PFAM terms), 103 new pathway predictions (KEGG-EC) and 96 new transmembrane domains (TMHMM terms). The updated annotation was assessed for completeness with BUSCO *v3*(38), and was shown to include a good representation of expected core eukaryotic genes, with minimal missing BUSCOs (2%) (Supplementary Figure 1).

Principal component analysis (PCA) of TMM normalised read counts per gene (Figure 2) showed that the biological replicates grouped closely together, with time points grouping into 4 clusters separated by time (PC1) and stage (PC2), as determined by K means clustering (Supplementary Figure 2).

**Figure 2:**
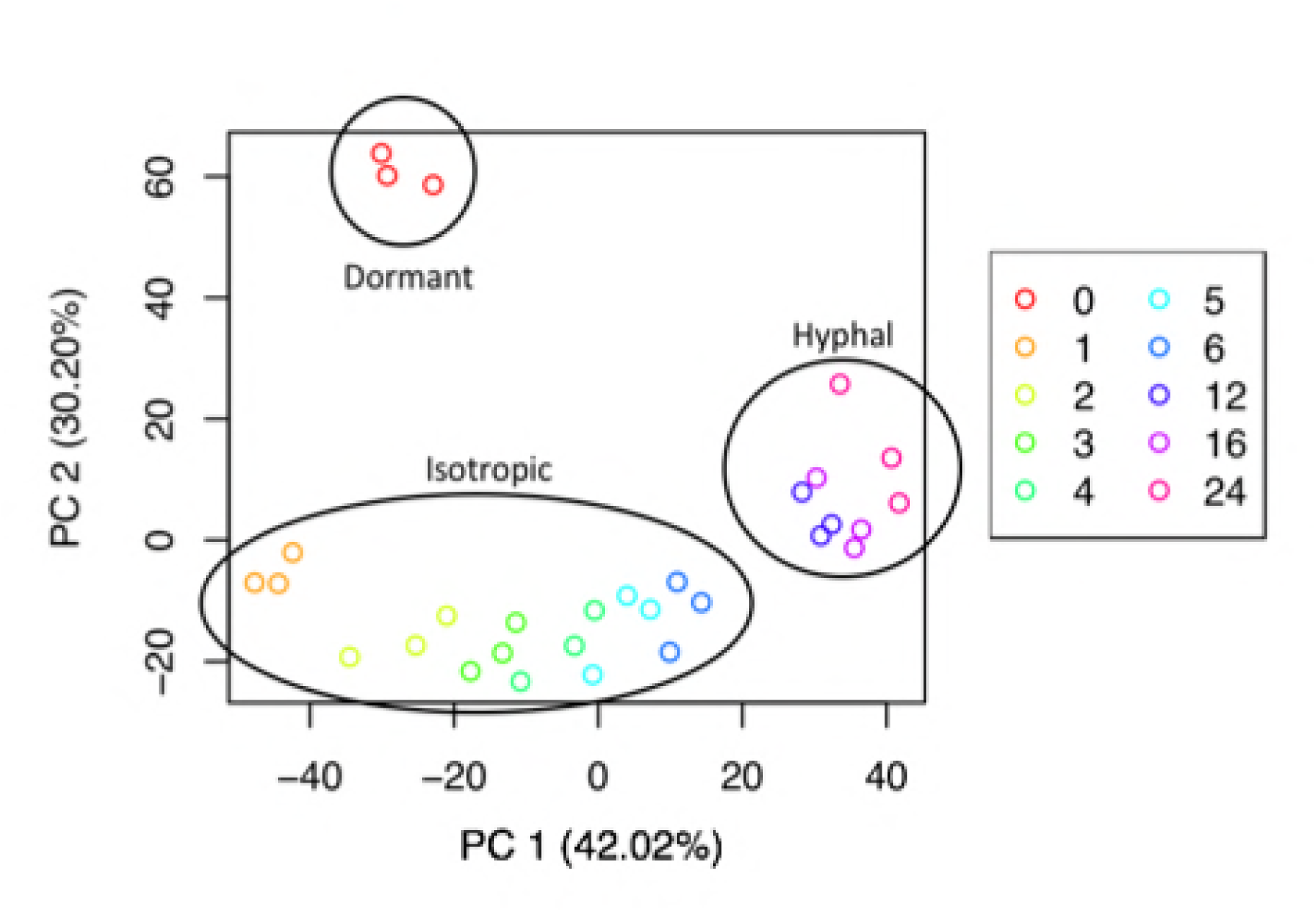
Principal component analysis of 7942 genes differentially expressed across all time points (n=3 for each time point. T= 0, 1,2,3,4,5,6,12,16,24 hours post germination). Each time point is color-coded.

### Dormant Spores are Transcriptionally Unique

In examining the overall transcriptional profiles of our cells, we observed a set of 482 transcripts that were only expressed in ungerminated spores (Figure 3A, T0), representing 3.76% of total transcripts expressed in ungerminated spores (Figure 3). As a result, genes expressed in resting spores account for 71.5% of all genes in the genome whereas the highest percentage of the genome covered by germinated spores is 68.8% (Figure 3A, T24). Resting spore specific transcripts that were co-expressed with other resting spore specific transcripts have predicted roles in lipid storage and localization, as well as transferase activity on phosphorous containing compounds (Figure 3B). As these transcripts are absent in germinated spores, they may have roles in the maintenance of spore dormancy.

**Figure 3:**
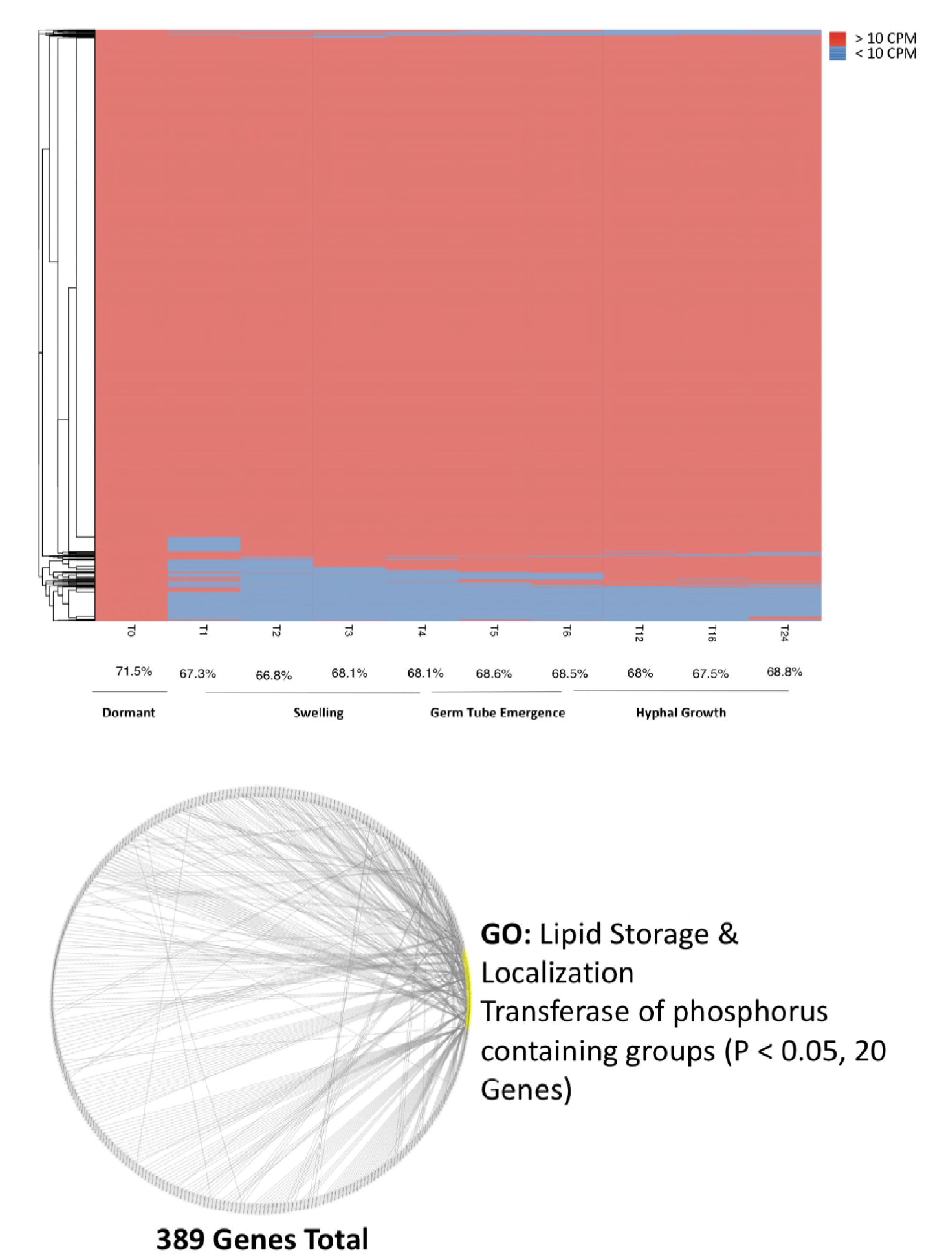
Resting spore specific expression. A) Heatmap displaying the absence (blue) or presence (red) of ten or more transcripts for a given gene over time. The average percent of the transcriptome expressed at any given time point is given below. B) Co-expression diagram, where each node represents a gene only expressed in ungerminated spores. Nodes linked to 10 or more others are highlighted in yellow, with their functions shown adjacent.

### Clustering of Transcriptional Changes Over Time

We performed a series of analyses to identify the transcriptional changes occurring during spore germination (Methods). PC analysis highlighted that the fungal transcriptome displayed a time dependent shift across 3 major clusters corresponding to the phenotypic developmental stages, swelling, germ tube emergence, and hyphal growth, indicating that spore germination is underpinned by progressive shifts in transcriptional regulation (Figure 2). The transcriptome of resting spores was distinct from that of all other developmental stages, changing dramatically between 0 and 1 hours. Thereafter, the transcriptional profiles of swelling spores and of those developing germ tubes were distinct but clustered together (2 to 6 hours). Furthermore, fully established filamentous growth was characterized by a specific transcriptional signature (12, 16 and 24 hours) (Figure 2). Consistent with stage specific transcriptional changes, progressive change in differential gene expression was observed during examination of the transcriptional profiles of each time point. A total of 7924 genes were differentially expressed across the entire time course (Figure 4A).

**Figure 4:**
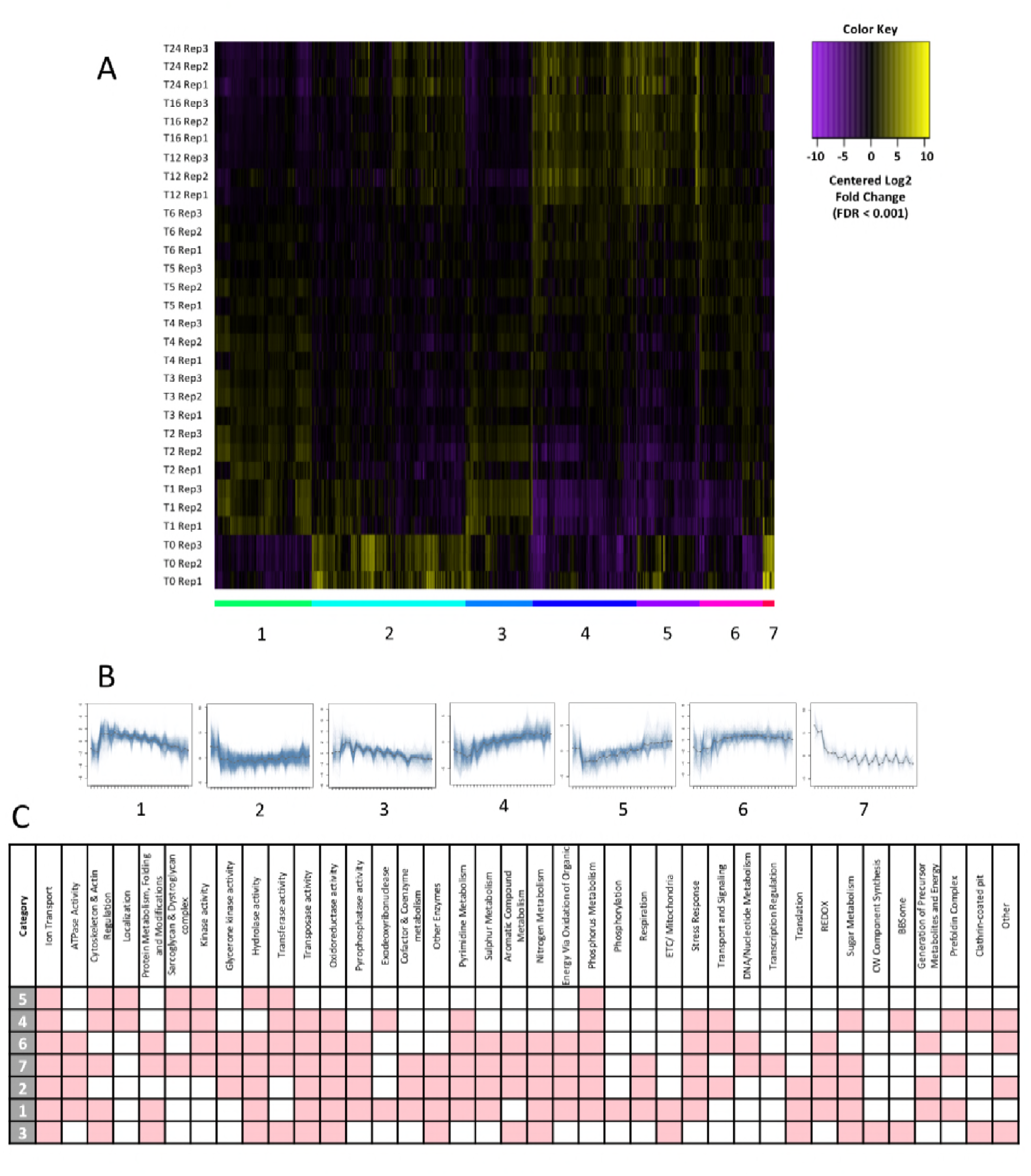
Clustering of expression over time. A) Heatmap displaying differentially expressed genes. Expression levels are plotted in Log2, space and mean-centered (FDR < 0.001) across the entire time course. K-means clustering has partitioned genes into 7 clusters, as indicated by coloured bars and numbered graphs below the heatmap. B) Graphs displaying cluster expression over time (0-24 hours). C) Table displaying categories enriched (Hypergeometric test, corrected P value < 0.05), indicated in red, for clusters 1 to 7.

Analysis of differentially expressed genes by k-means clustering identified seven major clusters of expression variation over time (Figure 4). Genes in clusters 1 and 3 are expressed at low levels in resting spores, with abundance increasing upon germination (1 hour; Figure 4B). Both clusters are enriched (Hypergeometric test, corrected P value < 0.05) for transcripts with predicted roles in regulation of the cytoskeleton, protein metabolism, the electron transport chain, translation and sugar metabolism (Figure 4C; Supplementary Table 1), suggesting these processes are important for germination initiation. Clusters 4 and 6 show gene expression levels moving from low to high over time, peaking during hyphal growth (Figure 4B). These clusters are enriched (Hypergeometric test, corrected P value < 0.05) for transcripts with predicted functions related to kinase, transferase, transposase and oxidoreductase activity, along with pyrimidine and phosphorous metabolism, stress response, transport and signalling (Figure 4C; Supplementary Table 1). This is consistent with the established roles for these processes in starting and maintaining vegetative growth(28, 39–41). Cluster 5 contains genes that have high expression levels in both ungerminated spores and the hyphal form, but low levels during initial swelling (Figure 4B). Cluster 5 is enriched (Hypergeometric test, corrected P value < 0.05) for transcripts with predicted functions in regulation of the cytoskeleton, transferase and hydrolase activity, and phosphorous metabolism (Figure 4C; Supplementary Table 1). This suggests that these functions may be repressed during isotropic growth to maintain swelling. Clusters 7 and 2 contain genes with expression levels peaking in ungerminated spores (Figure 4B). These clusters are enriched (Hypergeometric test, corrected P value < 0.05) for transcripts with predicted functions relating to glycerone kinase, pyrophosphatase, transferase, hydrolase and oxidoreductase activity, as well as cofactor and coenzyme metabolism, pyrimidine, sulphur, nitrogen, sugar and aromatic compound metabolism. These clusters are also enriched for reduction-oxidation (REDOX) processes, respiration and stress responses (Figure 4C; Supplementary Table 1). Notably, every cluster is enriched for transcripts involved in ion transport regulation, specifically potassium, sodium and hydrogen ions. This suggests tight regulation of transmembrane transport of these particular ions is important for the survival of *R. delemar*.

### Pairwise comparison of transcriptional changes over time correspond to phenotypic changes during germination

Ungerminated spores have a radically different expression profile to germinated spores (6456 significantly differentially expressed genes, FDR < 0.001), this is reflected by the functions of transcripts enriched in ungerminated spores. By pairwise comparisons of differentially expressed genes between time points, the largest transcriptional changes were seen during the first hour of germination (3,476 genes upregulated and 2,573 genes downregulated; Figure 5A). This was followed by a period of transcriptional consistency over the course of isotropic swelling, where few or no genes were found differentially expressed (Figure 5A). A noticeable shift in differential expression then bridges the beginning and later stages of hyphal growth (6-12 hours; Figure 5A). At the beginning of germination, an increase is observed in expression of transcripts with predicted roles in stress response, mitochondrial ribonucleases (MRP), the prefoldin complexes, organophosphate and sulphur metabolism, transposase, ATPase, nucleoside-triphosphatase and glycerone kinase activity (Figure 5B). A decrease in expression of genes with predicted functions in the organization of the actin cytoskeleton, carbohydrate metabolism, translation initiation factors, hexon binding, phosphodiesterase, arylformamidase, galactosylceramidase and precorrin-2 dehydrogenase activity is also seen (Figure 5B). Notably, some categories are both positively and negatively regulated at the beginning of germination: transcripts predicted to have roles in ion channel activity, hydrolase and pyrophosphatase activity do not always trend together (Figure 5B). It is likely these processes may involve several regulatory mechanisms implicated in initializing germination.

**Figure 5:**
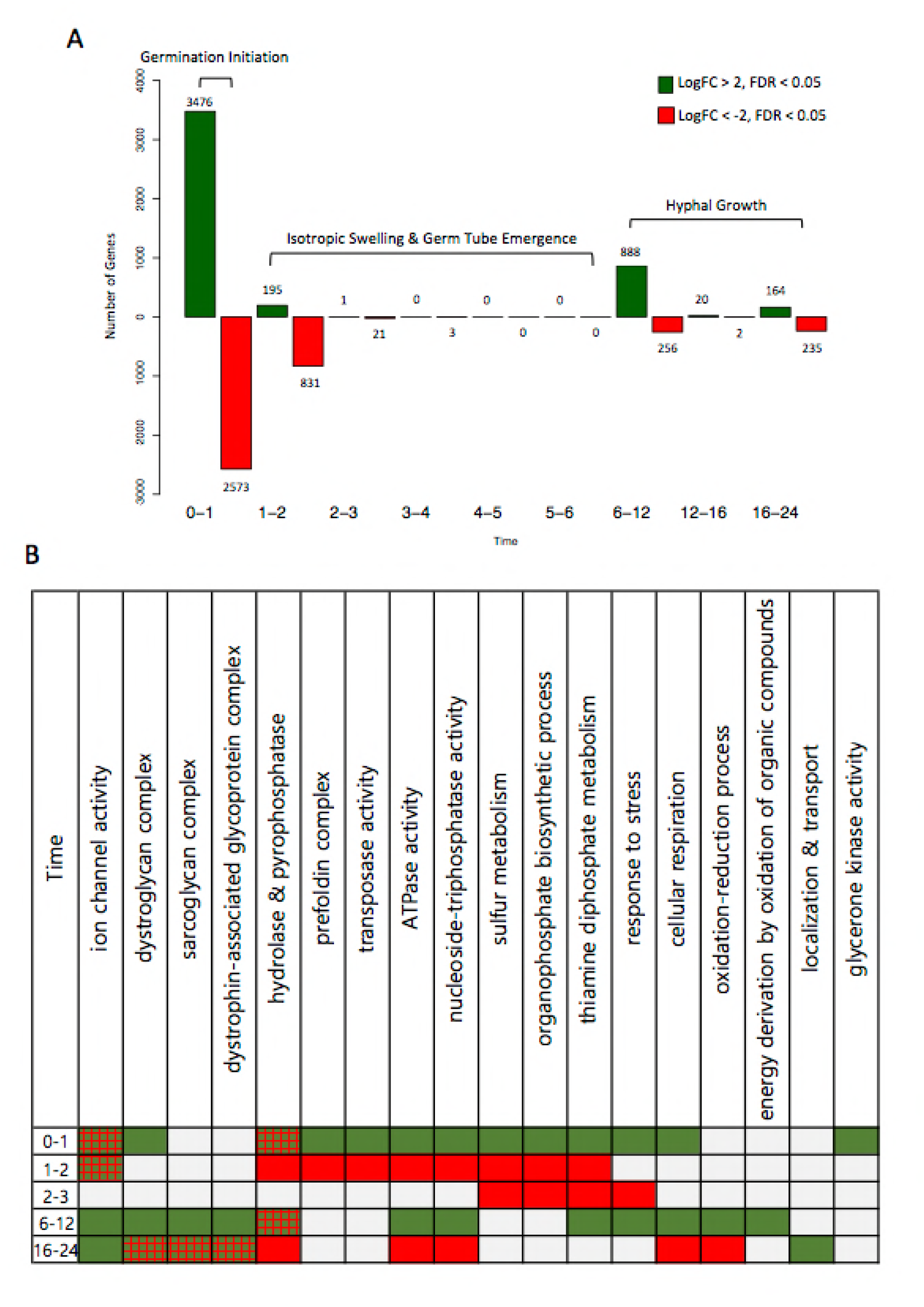
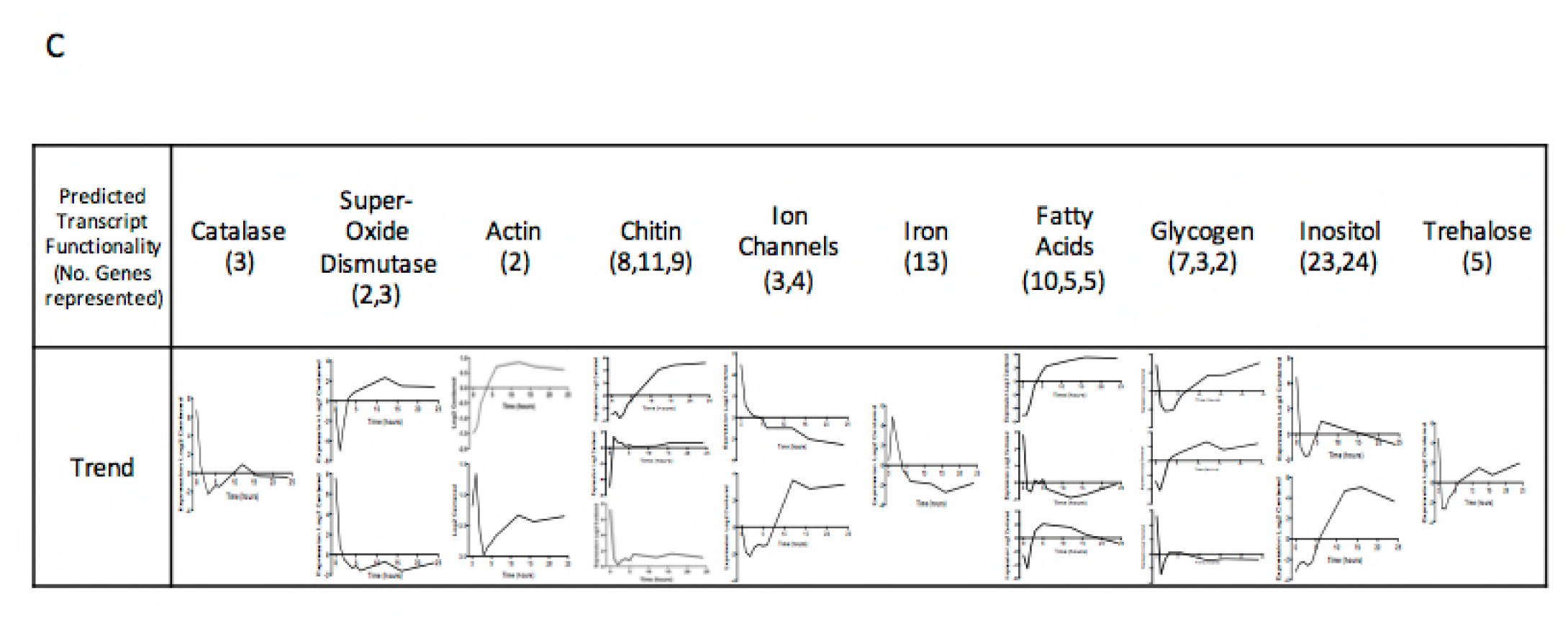
Differential gene expression over time A) The number of genes significantly differentially expressed (multiple corrected P value < 0.05) between time points, shown over time. Green bars indicate genes with an increase in expression (LogFC > 2), whilst red bars indicate genes with a decrease in expression (LogFC < -2). B) Enriched categories for the up/down regulated genes over time. Green boxes indicate an overall upregulation of this category, red indicates an overall down regulation and a red-green hatch indicates mixed regulation of this category. C) Expression profiles transcripts in specific categories over time, with the number of transcripts represented by each trend shown in brackets.

After initiation (1 - 2 hours), there is an overall trend of down regulation. The majority of transcripts that were upregulated at 1 hour are downregulated at 2 hours (Figure 5B), suggesting a reorganization of the transcriptome upon germination initiation. Notably, sulphur, organophosphate and thiamine diphosphate metabolism remain down regulated at both 2 and 3 hours. After the transcriptional stability during isotropic growth and hyphal emergence, transcripts with predicted roles in stress response, respiration, ATPase and nucleoside-triphosphatase activity and REDOX increase during early hyphal growth (Figure 5B). Between 6 and 12 hours, the proportion of down regulated transcripts decreases, with hydrolase and pyrophosphatase activity appearing both up and down regulated.

By examining expression profiles of predicted genes with biologically interesting functions (Figure 5C), we observe that iron acquisition transcripts rapidly increase during the initial phase of germination. This is consistent with literature which suggests iron scarcity induces abnormal germination and growth phenotypes in Mucorales species(42). Expression profiles for classes of genes related to actin, chitin and ion channels showed two or more contrasting trends (i.e., genes with the same class do not always travel together). However, when the opposing profiles are viewed simultaneously, we see upregulation in both ungerminated spores and the hyphal form. Phenotypic data indicates that the availability of chitin (calcofluor white stain (CFW); Figure 6A) within the cell wall increases rapidly over time, with spore cell walls containing high levels by 3 hours. The increase in cell wall protein content, denoted by fluorescein (FITC) staining (Figure 6A), also increases over time, with high concentrations present by 6 hours. Transcripts involved in the production and activity of trehalose, known as a stress response molecule in fungi(43), are also high in resting spores, but decrease upon initiation of germination. Consistent with a primed stress response, we observed that the ROS effectors SOD (Cu/Zn and Fe/Mn Super Oxide Dismutase) and catalase have increased expression levels in resting spores. These levels then decrease once germination is initiated, suggesting that a protective ROS stress response is involved in germination, perhaps to internal ROS produced through metabolic activity. We measured the production of endogenous ROS over time during germination (Figure 6A). We observed that the level of endogenously generated reactive oxygen species (ROS) within spores increases over the course of germination, but was limited to the spore body following germ tube emergence. We investigated the significance of ROS detoxification during germination by testing for resistance to exogenous (H_2_O_2_) and endogenous (mitochondrial derived) ROS (Figure 6B). Treatment with 5mM but not 1 mM H_2_O_2_ was sufficient to inhibit spore germination. In contrast, spores were highly sensitive to treatment with 1.5 or 10 nM Antimycin A, a mitochondrial inhibitor that impairs cytochrome C reductase activity leading to the accumulation of superoxide radicals within the cell. The impact of Antimycin A on germination may be two-fold, as we also observed that the expression of storage molecule transcripts appears high in both ungerminated spores and the hyphal form. High sensitivity to inhibition of oxidative phosphorylation with Antimycin A is consistent with reports that utilization of these storage molecules as energy reserves is important for the initiation and maintenance of growth(44–46).

**Figure 6:**
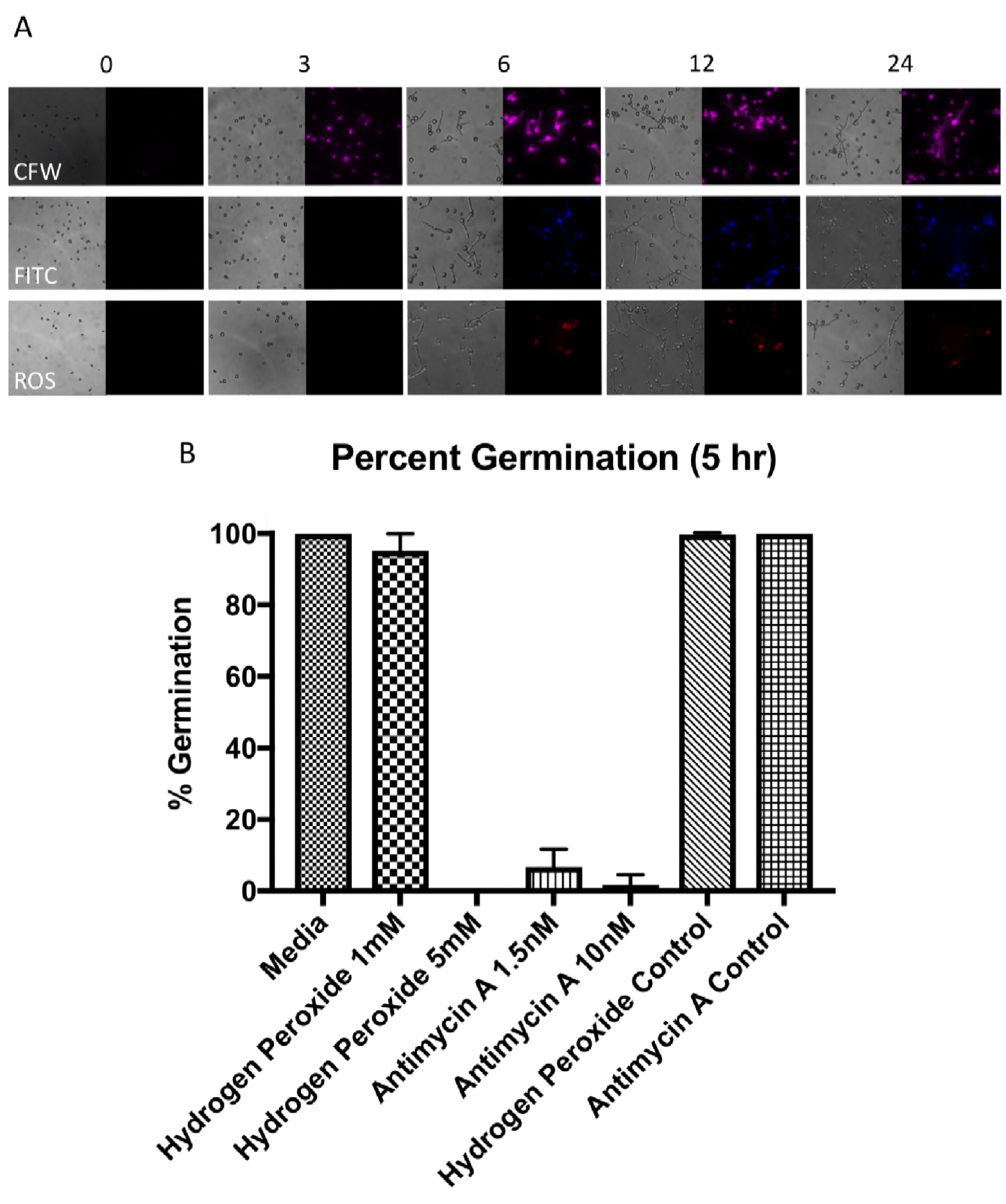
Cell wall dynamics and inhibition of germination. A) Spores germinated for 0,3,6,12 and 24 hours, stained with calcofluor white (CFW), fluorescein (FITC) and ROS stain carboxy-H_2_DCFDA (ROS). B) Germination is inhibited by 5mM of Hydrogen Peroxide and over 1.5nM of Antimycin A, as determined by live cell imaging, after 5 hours’ germination in SAB. Hydrogen peroxide control consists of an equivalent volume of H_2_0, Antimycin A control consists of an equivalent volume of 100% ethanol.

### Transcriptional hallmarks of germination are conserved across species, whilst R. delemar exhibits unique germination responses lacking in Aspergillus niger

It is unclear whether the mechanisms which underpin germination are conserved throughout the diverse fungal kingdom. To explore the extent of conservation, we compared our data set to other available transcriptional data sets for *Aspergillus niger* (Methods). When expression profiles of homologous genes from *A. niger* and *R. delemar* are compared over the course of germination, genes with common or unique functions specific to that time point can be identified. The largest shift in the transcriptional landscape of *A. niger* can be seen at the initial stage of germination(26, 28), we also observed this shift in *R. delemar* (Figure 7). Transcripts with predicted functions involved in transport and localization, proteolysis, glucose, hexose and carbohydrate metabolism increase at the initial stages of germination in both *A. niger* and *R. delemar*, whilst transcripts with predicted functions in translation, tRNA and rRNA processing, amine carboxylic acid and organic acid metabolism decrease. We also observe differences between the two datasets: over isotropic and hyphal growth, homologous genes with predicted functions in valine and branched chain amino acid metabolism were upregulated only in *R. delemar*, whilst homologous genes with predicted roles in ncRNA metabolism, translation, amino acid activation and ribosome biogenesis were downregulated exclusively in *R. delemar.* A 5% increase in genes which are uniquely up/down regulated in *R. delemar* are found in high synteny regions of the genome, when compared to genes which are up/down regulated in both *R. delemar* and *A. niger*. The duplicated nature of the *R. delemar* genome may allow for specific and tight regulation of the germination process, a feature unique to *R. delemar.*

**Figure 7:**
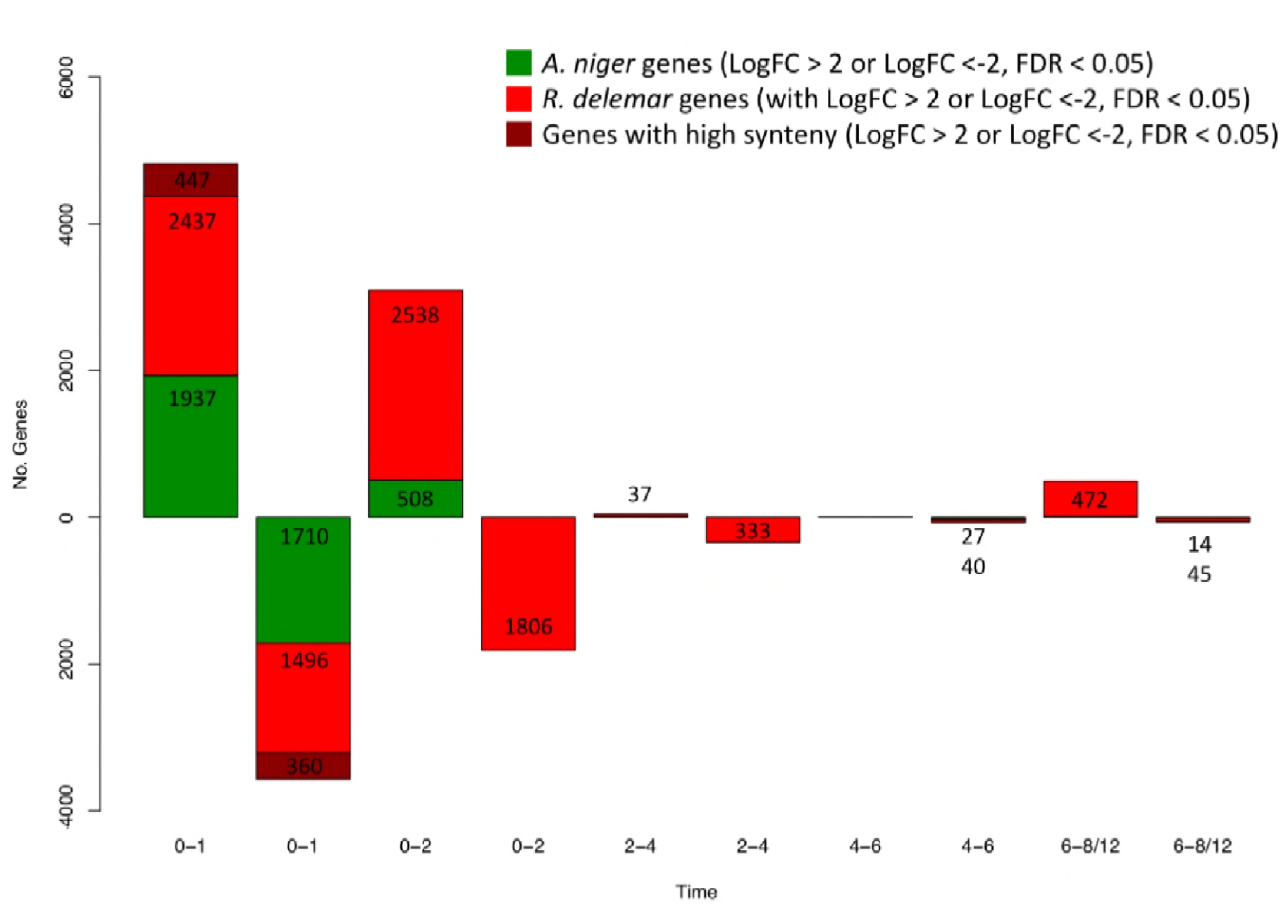
Number of homologues genes significantly differentially expressed (multiple corrected P value < 0.05) between time points, shown over time. Green represents the number of *A. niger* genes, red represents the number of *R. delemar* genes and dark red represents the number of *R. delemar* genes found in high synteny regions of the *R. delemar* genome.

## Discussion

Regulation of germination in the mucorales remains an under-examined area. Cues for germination include the availability of sufficient water, iron, a suitable carbon source and pH(10, 42, 47), though the mechanisms remain unclear. This study aims to expand current knowledge on the molecular processes which determine germination.

### Dormant Spores

Ungerminated spores show the least exposure of chitin and protein in the cell wall, suggesting these constituents may be masked prior to germination. It is established that the ungerminated conidia of various *Aspergillus* species are coated by a layer of hydrophobins which confer hydrophobicity to the conidia, and these structures rearrange upon germination to reveal a more heterogeneous and hydrophilic surface (48, 49). It may be the case that similar structures coat the outside of *R. delemar* spores prior to germination, inhibiting the visualisation of internal compounds such as chitin and protein. Transcripts involved in chitin processes, such as the predicted chitinases (Figure 5C), appear at higher levels in ungerminated spores, a feature which can also be seen in the dormant spores of *Aspergillus niger* (28). This suggests that the turnover or degradation of the fungal cell wall may be an important process involved in the formation of the spore, the maintenance of dormancy or the initial stages of germination. Pyrophosphatase, transferase, hydrolase and oxidoreductase activity also appear to be important in ungerminated spores. The presence of pyrophosphates has been implicated in aiding pathogenicity and survival in nutrient scarce environments for the fungal pathogen *Cryptococcus neoformans*(50). Interestingly, the signalling properties of pyrophosphates combined with inositol, also upregulated in ungerminated *R. delemar* spores, have been associated with metabolic regulation of yeast(51) and stress tolerance(52). There also appears to be a conserved requirement for sulphur in the early stages of germination across fungal species: sulphur and aromatic compound metabolism is upregulated in ungerminated spores of *R. delemar*, whilst sulphur metabolism is induced minutes after germination initiation in *Phomopsis viticola*. Sulphur has also been shown to be important for pathogenicity and the regulation of iron homeostasis in *A. fumigatus*(53). This may help explain the sharp increase in the levels of transcripts with predicted functions in iron recruitment upon the initiation of germination in *R. delemar* (Figure 5C). Resting spores also show an upregulation of transcripts involved in the latter stages of iron-sulphur cluster biosynthesis (Supplementary Figure 3).

Ungerminated spores are also enriched with transcripts involved in nitrogen metabolism. Nitrogen containing compounds have been shown to trigger germination in *A. niger*, correlating with the up regulation of transcripts involved in nitrogen utilisation during the initial stages of germination(28, 54).

Ungerminated *R. delemar* spores were also enriched for transcripts with roles in REDOX processes, respiration and stress responses. Predicted catalase, Cu/Zn and Fe/Mn superoxide dismutase genes appeared highly expressed in ungerminated spores, suggesting that they may form part of the stress response, as they are often utilised to resist internal metabolic ROS, as well as harsh conditions(55). An increased level of transcripts with predicted functions in the synthesis and phosphorylation of the stress response molecule trehalose(43) were also found in ungerminated spores (Figure 4C). This suggests regulation of trehalose processes may also be implicated in the resistance to harsh conditions by *R. delemar* spores.

Interestingly, transcripts only present in the ungerminated spores of *R. delemar* had roles in lipid storage. Lipid droplets have been observed in the spores of *Schizosaccharomyces pombe*, where it is thought they serve as energy reserves in nutritionally poor environments(56). It is likely these transcripts play roles in maintaining lipid storage molecules, crucial for spore survival in nutritionally scarce environments. Other transcripts unique to ungerminated spores had predicted roles in transference of phosphorous groups. Transcripts involved in the degradation of the phosphorous storage molecule phytate also appeared to be upregulated in ungerminated *R. delemar* spores, but downregulated upon the onset of germination. This indicates spores may depend on phosphorous reserves for the initiation of germination.

### Swelling Spores

During isotropic growth, the available chitin, protein and spore ROS content increases, and this is reflected by changes in the transcriptome. Transcripts predicted to play roles in cell wall biogenesis are enriched in cluster 3 (Figure 4), which shows an immediate increase in expression levels upon initiation of isotropic growth. Transcripts with roles in protein synthesis and modification are also enriched in cluster 3, amongst others. The observation that alterations in the structure and composition of the cell wall are seemingly required for germination suggests that identification of potential methods of inhibiting germination and therefore invasive infection is possible. For example, treatment with inhibitors of chitin synthesis and transporter machinery might offer a solution for inhibiting isotropic growth and germination(57). Predicted ROS scavenger transcripts such as catalase and some SODs are also downregulated after germination is initiated (Figure 5C). This correlates with the observation of increased levels of ROS in germinated spores. There does appear however to be a separate subset of Cu/Zn and Fe/Mn SOD transcripts which also remain abundant over time (Figure 5C, Upper Panel), providing a possible explanation as to why the swollen and hyphal forms are able to withstand the increased levels of ROS internally. ROS and SOD activity may also be involved in directing hyphal growth(40), or they may serve as signalling or metabolic molecules through compartmentalisation (58).

Clusters with expression levels that increase as isotropic growth begins (Figure 4) are also enriched in transcripts with predicted roles in the electron transport chain, translation and sugar metabolism. This suggests that respiration is a key metabolic process utilised throughout isotropic growth. Phenotypic data showing the inhibition of germination with Antimycin A also suggests this (Figure 6B). Protein synthesis is also required to manufacture new cellular machinery and prepare for hyphal emergence. *A. niger* also shows an increase in the production of transcripts involved in translation and respiration upon germination of conidia (28). Again, common themes in germination involving major category classes appear conserved throughout multiple families of filamentous fungi.

After the initial transcriptional shift, a higher proportion of transcripts are then down regulated by 2 hours (Figure 5). As the down regulated transcripts are mainly those that were upregulated at 1 hour, this ‘down regulation’ may be an artefact, as transcripts potentially essential for the initiation of germination are turned over or degraded following their use. Similarly, *A. niger* shows a vast downregulation of transcripts between 1 and 2 hours post initiation, though whether the majority of down regulated transcripts at this time point in *A. niger* are found in those upregulated at 1 hour has not been explored(26).

Notably, 17adiate, organophosphate and thiamine diphosphate metabolism are down regulated for initial and mid isotropic growth. Again, 17adiate utilization in *A. niger* appears under represented when transcripts from conidia having germinated for 2 hours are compared to those found in ungerminated conidia(28).

### Hyphal Growth

Hyphal samples were enriched for transcripts with predicted functions in kinase and oxidoreductase activity, as well as stress response, pyrimidine and phosphorous metabolism. Oxidoreductase is commonly used by the hyphal forms of wood decaying filamentous fungi such as *Phlebia 18adiate* and *Trichaptum abietinum*, thought to be useful for lignin decay(59). *R. delemar* is known to grow on plants with complex carbon sources(60), and the increased production of oxidoreductase may allow for the degradation of a variety of carbon sources, thus enabling *R. delemar* to colonise a variety of environments.

ROS levels appear to peak in the swollen bodies of hyphal R. *delemar*, whilst levels of transcripts with predicted functions in stress response also increase. Stress response genes have been shown to be important for the hyphal growth of the filamentous fungal plant pathogens *Fusarium graminearum* and *Ustilaginoidea virens*(61, 62). Furthermore, harsh environmental conditions can induce the production of ROS internally. For example, changes in osmolarity induce hydrogen peroxide bursts within the hyphae of *F. graminearum*(63). This remains to be studied in *Mucor* species. One of the central oxidative stress response transcription factors, Yap1, is found in a range of fungi including *Candida albicans, Aspergillus fumigatus* and *N. crassa,* and is essential for responding to ROS stress. When knocked out in *Epichloë festucae*, hyphae are susceptible to ROS(64). Unexpectedly a *YAP1* homologue could not be found in the genome of *R. delemar*. Together, our data highlight a role for ROS stress response in *R. delemar* germination and hyphal growth and suggest differences with other more well studied filamentous fungi.

During hyphal growth, functions enriched also included regulation of the cytoskeleton and phosphorous metabolism. The cytoskeleton is known to be important for hyphal extension, allowing the transport of vesicles to the hyphal tip to attain and maintain polarity(65, 66). Though phosphorous metabolism in the hyphae of filamentous fungi is not as well studied, it has been shown that phosphorus levels in the soil can effect germination and hyphal extension length in mycorrhizal fungi(67).

As hypothesised, transcripts with predicted roles in respiration also appear to peak around hyphal growth in *R. delemar*. This appears to be a conserved trait across filamentous fungi, as a higher respiratory rate is commonly seen in the hyphal form of *Trichoderma lignorum*(68), and increased levels of respiratory transcripts are present in hyphae of *N. crassa*(21).

The results of this study increase our understanding of the molecular mechanisms controlling germination in *R. delemar*. We have shown that ungerminated spores are transcriptionally unique, whilst the initiation of germination entails a huge transcriptional shift. ROS resistance and respiration are required for germination to occur, whilst actin, chitin and cytoskeletal components appear to play key roles initiating isotropic swelling and hyphal growth. *R. delemar* shares many transcriptional traits with *A. niger* at germination initiation, however unique transcriptional features to *R. delemar* indicate that the duplicated nature of the genome may allow for alternative regulation of this process. This study has provided a significant overview of the transcriptome of germinating spores and expanded current knowledge in the mucorales field.

## Materials and Methods

### Culture

*R. delemar* was cultured with Sabouraud dextrose agar or broth (10 g/L mycological peptone, 20 g/L dextrose) sourced from Sigma-Aldrich, at room temperature. Spores were harvested with PBS, centrifuged for 3 minutes at 3000 rpm and washed. Appropriate concentrations of spores were used for further experiments.

### Live Cell Imaging, Staining and Inhibition

Images of 1×10^5^Spores/ml in SAB broth were taken every 10 minutes to determine germination characteristics. Images were taken at 20X objective on a Zeiss Axio Observer. Calcofluor white (CFW), Fluorescein (FITC) (Sigma-Aldrich) and the ROS stain carboxy-H_2_DCFDA (C400) (Invitrogen) were incubated with live spores, according to manufacturer’s instructions, prior to imaging. To assess inhibition, spores were incubated with 1-5mM of Hydrogen peroxide, or 1.5-10nM of Antimycin A (Sigma-Aldrich) prior to imaging. Bright field and fluorescent images were then analysed using ImageJ V1.

### RNA Extraction and Sequencing

Total RNA was extracted from *R. delemar* spores which germinated in SAB broth for 0,1,2,3,4,5,6,12,16 and 24 hours. To extract total RNA, the washed samples were immediately immersed in Trizol and lysed via bead beating at 6500 rpm for 60 seconds. Samples were then either immediately frozen at -20°C and stored for RNA extraction, or placed on ice for RNA extraction. After lysis, 0.2ml of chloroform was added for every 1ml of Trizol used in the sample preparation. Samples were incubated for 3 minutes, then spun at 12,000 G at 4°C for 15 minutes. To the aqueous phase, an equal volume of 100% EtOH was added, before the samples were loaded onto RNeasy RNA extraction columns (Qiagen). Manufacturer’s instructions were followed from this point onwards. RNA quality was checked by Agilent, with all RIN scores above 8 (69). 1ug of Total RNA was used for cDNA library preparation. Library preparation was done in accordance with the NEBNext pipeline, with library quality checked by Agilent. Samples were sequenced using the Illumina HiSeq platform, 100bp paired end sequencing was employed (2×100bp).

### Data Analysis

Fastqc (version 0.11.5) was employed to ensure the quality of all samples, a Phred value of over 36 was found for every sample. Hisat2 (version 2.0.5) was used to align reads to the indexed genome of *Rhizopus delemar* found on JGI (PRJNA13066)(35, 70) HTSeq (version 0.8.0) was used to quantify the output(71). Trinity and edgeR (version 3.16.5) were then used to analyse differential expression(72, 73). Pathway tools (version 21.0) was used to obtain information on specific pathways(74). The Genome of *R. delemar* was re-annotated by incorporating the RNA-Seq data via Braker (version 2.1.0), this was fed into the Broad institute annotation pipeline, which removed sequences with overlapped with repetitive elements, numbered, and named genes as previously described(75). Completeness of annotation was analysed with BUSCO (version 3)(37, 38).

### Accession numbers

Raw data and a compiled count matrix can be obtained with the following accession numbers: *SRP146252* (SRA).

## Acknowledgments

The authors would like to thank the fungal genetic stock centre for providing the *Rhizopus delemar* strain RA 99-880 and the University of Birmingham Genomic Services facility for conducting the sequencing. PSC was funded by BBSRC MIBTP PhD studentship and a travel stipend from the Microbiological Society. ERB was funded by the BBSRC (BB/M014525/1). KV was funded by the Wellcome Trust (108387/Z/15/Z). JFM and CAC were funded by NIAID (U19AI110818) to the Broad Institute.

## Supplementary figure legends

Figure 1. Table displaying genome annotation statistics, comparing the original *R. delemar* annotation (column 1) and the annotation updated with our RNA-Seq data (column 2)

Figure 2. Sum of squares cluster analysis of PCA data

Figure 3. Upregulation (red) of the Fe-S cluster biosynthetic pathway, determined with PathwayTools

Table 1. Listing of enriched terms.

## Re Annotation of *R. delemar* Genome

**Figure 1:**
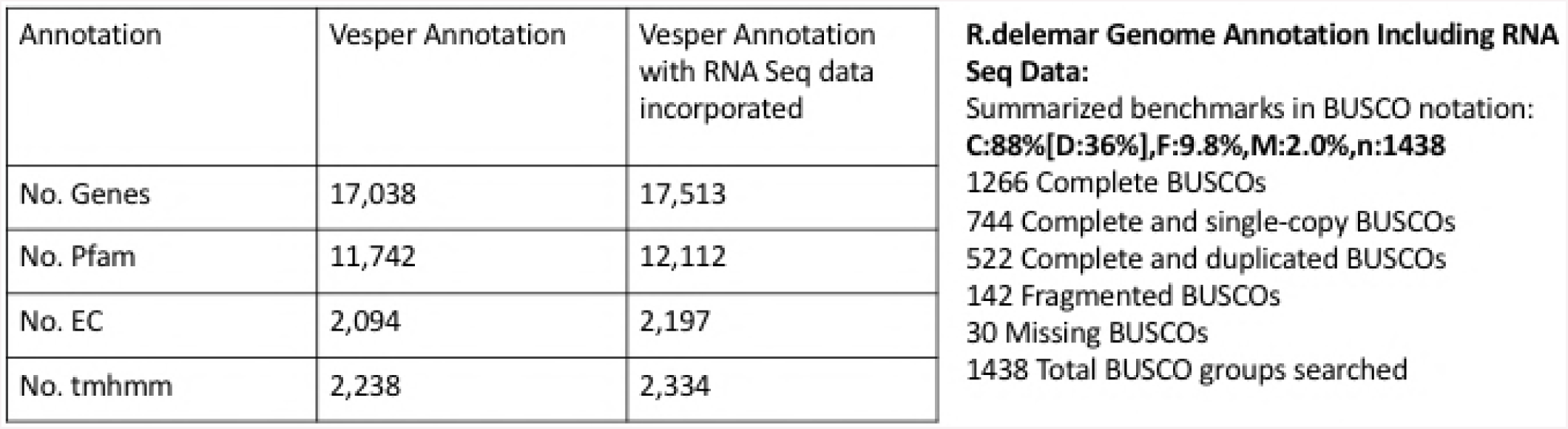
Table displaying genome annotation statistics, comparing the original R. delemar annotation {column 1) and the annotation updated with our RNA-Seq data (column 2)

**Figure 2:**
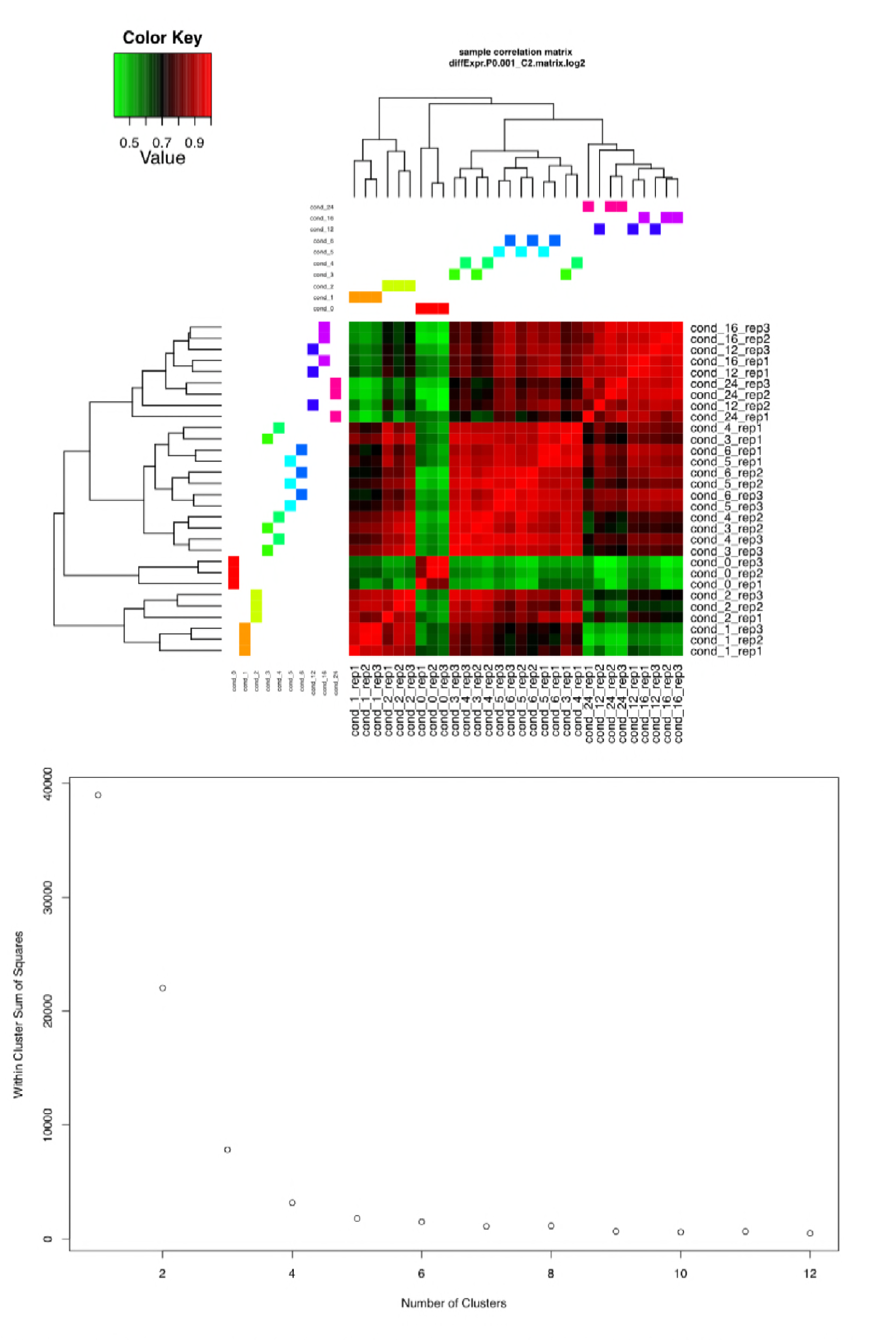
Sample correlation and sum of squares cluster analysis of principle component analysis data

**Figure 3:**
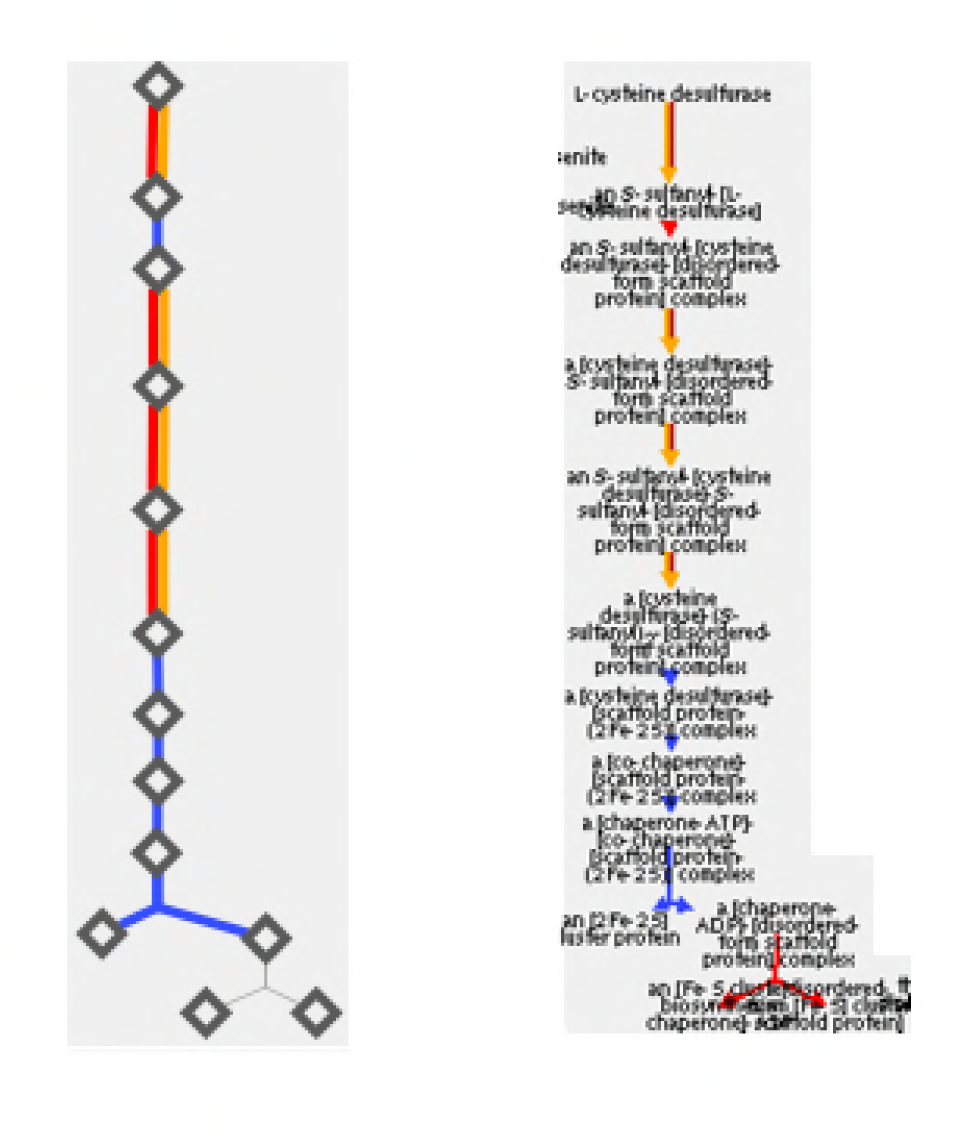
Upregulation (red} of the Fe-5 cluster biosynthetic pathway, determined with PathwayTools

~~~
Cluster 1
term_id pvalue adjp term_name
GO:0015459 6.60E-16 4.80E-14 potassium channel regulator activity
GO:0019870 6.60E-16 4.80E-14 potassium channel inhibitor activity
GO:0016248 1.70E-14 6.00E-13 channel inhibitor activity
GO:0008200 1.70E-14 6.00E-13 ion channel inhibitor activity
GO:0016247 2.20E-14 6.40E-13 channel regulator activity
GO:0042302 1.80E-05 0.00044 structural constituent of cuticle
GO:0044087 2.10E-06 0.00056 regulation of cellular component biogenesis
GO:0006298 1.80E-05 0.0012 mismatch repair
GO:0016310 3.00E-05 0.0012 phosphorylation
GO:0051694 3.40E-05 0.0012 pointed-end actin filament capping
GO:1901879 5.60E-05 0.0012 regulation of protein depolymerization
GO:0031333 5.60E-05 0.0012 negative regulation of protein complex assembly
GO:0030834 5.60E-05 0.0012 regulation of actin filament depolymerization
GO:0032272 5.60E-05 0.0012 negative regulation of protein polymerization
GO:0051494 5.60E-05 0.0012 negative regulation of cytoskeleton organization
GO:1901880 5.60E-05 0.0012 negative regulation of protein depolymerization
GO:0030835 5.60E-05 0.0012 negative regulation of actin filament depolymerization
GO:0030837 5.60E-05 0.0012 negative regulation of actin filament polymerization
GO:0051693 5.60E-05 0.0012 actin filament capping
GO:0033554 6.20E-05 0.0012 cellular response to stress
GO:0051302 7.90E-05 0.0014 regulation of cell division
GO:0016272 2.50E-05 0.0014 prefoldin complex
GO:0043242 9.00E-05 0.0015 negative regulation of protein complex disassembly
GO:0090066 0.00014 0.0015 regulation of anatomical structure size
GO:0032535 0.00014 0.0015 regulation of cellular component size
GO:0032970 0.00014 0.0015 regulation of actin filament-based process
GO:0043254 0.00014 0.0015 regulation of protein complex assembly
GO:0030832 0.00014 0.0015 regulation of actin filament length
GO:0032271 0.00014 0.0015 regulation of protein polymerization
GO:0032956 0.00014 0.0015 regulation of actin cytoskeleton organization
GO:0030833 0.00014 0.0015 regulation of actin filament polymerization
GO:0008064 0.00014 0.0015 regulation of actin polymerization or depolymerization
GO:0010639 2.00E-04 0.002 negative regulation of organelle organization
GO:0042357 0.00021 0.002 thiamine diphosphate metabolic process
GO:0009229 0.00021 0.002 thiamine diphosphate biosynthetic process
GO:0006950 0.00022 0.002 response to stress
GO:0044445 9.60E-05 0.002 cytosolic part
GO:0005829 1.00E-04 0.002 cytosol
GO:0006793 0.00024 0.0021 phosphorus metabolic process
GO:0015980 0.00027 0.0023 energy derivation by oxidation of organic compounds
GO:0006796 0.00031 0.0025 phosphate-containing compound metabolic process
GO:0032465 0.00033 0.0025 regulation of cytokinesis
GO:0046939 0.00033 0.0025 nucleotide phosphorylation
GO:0006165 0.00033 0.0025 nucleoside diphosphate phosphorylation
GO:0055114 0.00034 0.0025 oxidation-reduction process
GO:0006281 0.00037 0.0027 DNA repair
GO:0006091 0.00049 0.0034 generation of precursor metabolites and energy
GO:0000172 0.00026 0.0037 ribonuclease MRP complex
GO:0009132 0.00057 0.0039 nucleoside diphosphate metabolic process
GO:0044272 0.00062 0.004 sulfur compound biosynthetic process
GO:0006974 0.00062 0.004 response to DNA damage stimulus
GO:0043244 0.00068 0.0043 regulation of protein complex disassembly
GO:0016887 0.00021 0.0043 ATPase activity
GO:0051129 0.00078 0.0047 negative regulation of cellular component organization
GO:0051493 0.00078 0.0047 regulation of cytoskeleton organization
GO:0005732 0.00042 0.0048 small nucleolar ribonucleoprotein complex
GO:0008152 9.00E-04 0.0054 metabolic process
GO:0090407 0.001 0.0059 organophosphate biosynthetic process
GO:0044237 0.0013 0.0071 cellular metabolic process
GO:0006790 0.0013 0.0071 sulfur compound metabolic process
GO:0006811 0.0013 0.0071 ion transport
GO:0045333 0.0015 0.0078 cellular respiration
GO:0017111 0.00054 0.0096 nucleoside-triphosphatase activity
GO:0032991 0.0012 0.012 macromolecular complex
GO:0019637 0.0027 0.014 organophosphate metabolic process
GO:0072528 0.0029 0.015 pyrimidine-containing compound biosynthetic process
GO:0051128 0.0033 0.017 regulation of cellular component organization
GO:0044271 0.0033 0.017 cellular nitrogen compound biosynthetic process
GO:0005198 0.001 0.017 structural molecule activity
GO:0016462 0.0018 0.019 pyrophosphatase activity
GO:0004529 0.0019 0.019 exodeoxyribonuclease activity
GO:0016895 0.0019 0.019 “exodeoxyribonuclease activity, producing 5′-phosphomonoesters”
GO:0016818 0.002 0.019 “hydrolase activity, acting on acid anhydrides, in phosphorus-containing anhydrides”
GO:0008409 0.0022 0.019 5′-3′ exonuclease activity
GO:0035312 0.0022 0.019 5′-3′ exodeoxyribonuclease activity
GO:0008297 0.0022 0.019 single-stranded DNA specific exodeoxyribonuclease activity
GO:0045145 0.0022 0.019 single-stranded DNA specific 5′-3′ exodeoxyribonuclease activity
GO:0016817 0.0024 0.019 “hydrolase activity, acting on acid anhydrides”
GO:0004371 0.0025 0.019 glycerone kinase activity
GO:0009432 0.0044 0.022 SOS response
GO:0051716 0.0047 0.022 cellular response to stimulus
GO:0071855 0.003 0.022 neuropeptide receptor binding
GO:0004803 0.0033 0.023 transposase activity
GO:0033043 0.0051 0.024 regulation of organelle organization
GO:0022904 0.0051 0.024 respiratory electron transport chain
GO:0009308 0.0054 0.024 amine metabolic process
GO:0006006 0.0056 0.025 glucose metabolic process
GO:0051188 0.0057 0.025 cofactor biosynthetic process
GO:0031423 0.0039 0.025 hexon binding
GO:0005789 0.0034 0.028 endoplasmic reticulum membrane
GO:0042175 0.0043 0.028 nuclear outer membrane-endoplasmic reticulum membrane network
GO:0044432 0.0043 0.028 endoplasmic reticulum part
GO:0043234 0.0051 0.029 protein complex
GO:0005783 0.0055 0.029 endoplasmic reticulum
GO:0030686 0.0059 0.029 90S preribosome
GO:0051186 0.0078 0.034 cofactor metabolic process
GO:0072527 0.0093 0.039 pyrimidine-containing compound metabolic process
GO:0006835 0.01 0.043 dicarboxylic acid transport
GO:0019318 0.011 0.046 hexose metabolic process
GO:0048523 0.012 0.048 negative regulation of cellular process
GO:1901681 0.0076 0.048 sulfur compound binding
Cluster2
term_id pvalue adjp term_name
GO:0006298 9.20E-07 0.00016 mismatch repair
GO:0090407 1.00E-05 0.00092 organophosphate biosynthetic process
GO:0072528 2.30E-05 0.0011 pyrimidine-containing compound biosynthetic process
GO:0042357 3.20E-05 0.0011 thiamine diphosphate metabolic process
GO:0009229 3.20E-05 0.0011 thiamine diphosphate biosynthetic process
GO:0006281 5.20E-05 0.0014 DNA repair
GO:0018130 5.60E-05 0.0014 heterocycle biosynthetic process
GO:0019438 8.20E-05 0.0015 aromatic compound biosynthetic process
GO:0006974 8.40E-05 0.0015 response to DNA damage stimulus
GO:0044271 8.80E-05 0.0015 cellular nitrogen compound biosynthetic process
GO:0072527 9.20E-05 0.0015 pyrimidine-containing compound metabolic process
GO:1901362 0.00011 0.0016 organic cyclic compound biosynthetic process
GO:0019637 0.00038 0.0048 organophosphate metabolic process
GO:0006796 0.00038 0.0048 phosphate-containing compound metabolic process
GO:0044237 0.00043 0.0051 cellular metabolic process
GO:0032465 5.00E-04 0.0055 regulation of cytokinesis
GO:0006793 0.00099 0.0098 phosphorus metabolic process
GO:0044087 0.001 0.0098 regulation of cellular component biogenesis
GO:0071855 2.00E-04 0.0098 neuropeptide receptor binding
GO:0016247 0.00032 0.0098 channel regulator activity
GO:0015459 0.00035 0.0098 potassium channel regulator activity
GO:0019870 0.00035 0.0098 potassium channel inhibitor activity
GO:0051302 0.0011 0.01 regulation of cell division
GO:0006732 0.0013 0.012 coenzyme metabolic process
GO:0031423 0.00084 0.012 hexon binding
GO:0016248 0.00089 0.012 channel inhibitor activity
GO:0008200 0.00089 0.012 ion channel inhibitor activity
GO:0016887 0.001 0.012 ATPase activity
GO:0016462 0.0011 0.012 pyrophosphatase activity
GO:0004803 0.0012 0.012 transposase activity
GO:0016818 0.0012 0.012 “hydrolase activity, acting on acid anhydrides, in phosphorus-containing anhydrides”
GO:0017111 0.0013 0.012 nucleoside-triphosphatase activity
GO:0016817 0.0015 0.012 “hydrolase activity, acting on acid anhydrides”
GO:0015980 0.0017 0.013 energy derivation by oxidation of organic compounds
GO:0045333 0.0017 0.013 cellular respiration
GO:0007186 0.0017 0.013 G-protein coupled receptor signaling pathway
GO:0008152 0.0018 0.013 metabolic process
GO:0051186 0.0019 0.013 cofactor metabolic process
GO:0009108 0.0019 0.013 coenzyme biosynthetic process
GO:0051716 0.0021 0.014 cellular response to stimulus
GO:0005057 0.0018 0.014 receptor signaling protein activity
GO:0030515 0.0019 0.014 snoRNA binding
GO:0033554 0.0024 0.015 cellular response to stress
GO:0006091 0.0026 0.016 generation of precursor metabolites and energy
GO:0051188 0.0027 0.016 cofactor biosynthetic process
GO:0044272 0.0029 0.017 sulfur compound biosynthetic process
GO:0055114 0.003 0.017 oxidation-reduction process
GO:1901360 0.0039 0.021 organic cyclic compound metabolic process
GO:0046483 0.004 0.021 heterocycle metabolic process
GO:0006835 0.0045 0.023 dicarboxylic acid transport
GO:0006790 0.0048 0.023 sulfur compound metabolic process
GO:0006811 0.0048 0.023 ion transport
GO:0004784 0.0035 0.023 superoxide dismutase activity
GO:0016721 0.0035 0.023 “oxidoreductase activity, acting on superoxide radicals as acceptor”
GO:0006725 0.0055 0.026 cellular aromatic compound metabolic process
GO:0034641 0.0062 0.028 cellular nitrogen compound metabolic process
GO:0007166 0.0064 0.028 cell surface receptor signaling pathway
GO:0004371 0.0057 0.035 glycerone kinase activity
GO:1901566 0.011 0.049 organonitrogen compound biosynthetic process
Cluster 3
term_id pvalue adjp term_name
GO:0000313 9.80E-41 3.80E-39 organellar ribosome
GO:0005761 9.80E-41 3.80E-39 mitochondrial ribosome
GO:0005840 6.60E-40 1.70E-38 ribosome
GO:0005759 1.70E-39 3.20E-38 mitochondrial matrix
GO:0000270 6.00E-36 1.20E-33 peptidoglycan metabolic process
GO:0009252 6.70E-36 1.20E-33 peptidoglycan biosynthetic process
GO:0009273 1.70E-35 1.50E-33 peptidoglycan-based cell wall biogenesis
GO:0006024 1.70E-35 1.50E-33 glycosaminoglycan biosynthetic process
GO:0006023 4.10E-35 2.90E-33 aminoglycan biosynthetic process
GO:0044429 5.60E-34 8.70E-33 mitochondrial part
GO:0015020 1.60E-34 1.20E-32 glucuronosyltransferase activity
GO:0015018 1.60E-34 1.20E-32 galactosylgalactosylxylosylprotein 3-betaglucuronosyltransferase activity
GO:0030203 4.40E-34 2.40E-32 glycosaminoglycan metabolic process
GO:0070589 5.50E-34 2.40E-32 cellular component macromolecule biosynthetic process
GO:0044038 5.50E-34 2.40E-32 cell wall macromolecule biosynthetic process
GO:0005739 2.10E-33 2.70E-32 mitochondrion
GO:0044036 2.90E-33 1.10E-31 cell wall macromolecule metabolic process
GO:0042546 3.10E-32 1.10E-30 cell wall biogenesis
GO:0006022 1.20E-29 3.70E-28 aminoglycan metabolic process
GO:0043233 4.30E-28 4.20E-27 organelle lumen
GO:0070013 4.30E-28 4.20E-27 intracellular organelle lumen
GO:0031974 6.90E-28 6.00E-27 membrane-enclosed lumen
GO:0030529 2.40E-26 1.90E-25 ribonucleoprotein complex
GO:0071554 1.50E-26 4.30E-25 cell wall organization or biogenesis
GO:0008194 1.20E-26 6.40E-25 UDP-glycosyltransferase activity
GO:0034464 1.00E-25 7.10E-25 BBSome
GO:0044085 4.00E-24 1.10E-22 cellular component biogenesis
GO:0016758 5.50E-23 2.10E-21 “transferase activity, transferring hexosyl groups”
GO:1901137 1.50E-22 3.90E-21 carbohydrate derivative biosynthetic process
GO:0016757 1.80E-21 5.50E-20 “transferase activity, transferring glycosyl groups”
GO:0043228 8.30E-20 5.00E-19 non-membrane-bounded organelle
GO:0043232 8.30E-20 5.00E-19 intracellular non-membrane-bounded organelle
GO:0044422 3.10E-17 1.70E-16 organelle part
GO:0044446 3.30E-17 1.70E-16 intracellular organelle part
GO:0043231 9.20E-16 4.50E-15 intracellular membrane-bounded organelle
GO:0043227 1.10E-15 5.10E-15 membrane-bounded organelle
GO:1901135 3.40E-16 8.00E-15 carbohydrate derivative metabolic process
GO:1901566 7.80E-13 1.70E-11 organonitrogen compound biosynthetic process
GO:0043229 1.00E-11 4.40E-11 intracellular organelle
GO:0043226 1.30E-11 5.50E-11 organelle
GO:0034645 9.10E-12 1.90E-10 cellular macromolecule biosynthetic process
GO:0009059 8.50E-11 1.70E-09 macromolecule biosynthetic process
GO:1901564 4.10E-10 7.60E-09 organonitrogen compound metabolic process
GO:0071840 2.90E-09 5.10E-08 cellular component organization or biogenesis
GO:0031532 1.50E-08 2.60E-07 actin cytoskeleton reorganization
GO:0004553 8.10E-07 1.90E-05 “hydrolase activity, hydrolyzing O-glycosyl compounds”
GO:0004113 8.50E-07 1.90E-05 “2′,3′-cyclic-nucleotide 3′-phosphodiesterase activity”
GO:0004336 1.00E-06 2.00E-05 galactosylceramidase activity
GO:0044710 2.90E-06 4.60E-05 single-organism metabolic process
GO:0016798 3.50E-06 6.00E-05 “hydrolase activity, acting on glycosyl bonds”
GO:0005099 1.40E-05 0.00022 Ras GTPase activator activity
GO:0031423 2.80E-05 0.00039 hexon binding
GO:0004061 3.30E-05 0.00043 arylformamidase activity
GO:0044699 3.00E-05 0.00047 single-organism process
GO:1901576 3.90E-05 0.00058 organic substance biosynthetic process
GO:0030029 5.80E-05 0.00079 actin filament-based process
GO:0030036 5.80E-05 0.00079 actin cytoskeleton organization
GO:0044260 7.10E-05 0.00088 cellular macromolecule metabolic process
GO:0009058 7.30E-05 0.00088 biosynthetic process
GO:0051821 7.70E-05 0.00088 dissemination or transmission of organism from
other organism involved in symbiotic interaction
GO:0044007 7.70E-05 0.00088 dissemination or transmission of symbiont from host
GO:0019089 7.70E-05 0.00088 transmission of virus
GO:0044765 0.00011 0.0012 single-organism transport
GO:0044249 0.00012 0.0013 cellular biosynthetic process
GO:0009396 0.00021 0.0021 folic acid-containing compound biosynthetic process
GO:0006826 0.00021 0.0021 iron ion transport
GO:0043170 0.00022 0.0021 macromolecule metabolic process
GO:0016740 0.00018 0.0021 transferase activity
GO:0004556 0.00021 0.0024 alpha-amylase activity
GO:0009521 0.00067 0.0026 photosystem
GO:0016160 0.00025 0.0026 amylase activity
GO:0042559 0.00042 0.004 pteridine-containing compound biosynthetic process
GO:0048513 0.00046 0.0041 organ development
GO:0032312 0.00046 0.0041 regulation of ARF GTPase activity
GO:0006807 0.00052 0.0046 nitrogen compound metabolic process
GO:0040011 0.00057 0.0049 locomotion
GO:0031369 0.00064 0.0059 translation initiation factor binding
GO:0005096 0.00065 0.0059 GTPase activator activity
GO:0006760 0.00076 0.0064 folic acid-containing compound metabolic process
GO:0006811 0.00082 0.0067 ion transport
GO:0051234 0.00089 0.007 establishment of localization
GO:0006810 0.00089 0.007 transport
GO:0043648 0.00092 0.0071 dicarboxylic acid metabolic process
GO:0030705 0.0011 0.008 cytoskeleton-dependent intracellular transport
GO:0042073 0.0011 0.008 intraflagellar transport
GO:0010970 0.0011 0.008 microtubule-based transport
GO:0032012 0.0011 0.008 regulation of ARF protein signal transduction
GO:0019887 0.00093 0.0081 protein kinase regulator activity
GO:0019207 0.0011 0.009 kinase regulator activity
GO:0019210 0.0015 0.011 kinase inhibitor activity
GO:0004860 0.0015 0.011 protein kinase inhibitor activity
GO:0044782 0.0017 0.012 cilium organization
GO:0015459 0.0017 0.012 potassium channel regulator activity
GO:0019870 0.0017 0.012 potassium channel inhibitor activity
GO:0005083 0.0018 0.012 small GTPase regulator activity
GO:0043115 0.002 0.012 precorrin-2 dehydrogenase activity
GO:0048731 0.0024 0.016 system development
GO:0046907 0.0024 0.016 intracellular transport
GO:0007010 0.0027 0.018 cytoskeleton organization
GO:0016787 0.003 0.018 hydrolase activity
GO:0012505 0.0051 0.019 endomembrane system
GO:0004857 0.0035 0.019 enzyme inhibitor activity
GO:0030695 0.0035 0.019 GTPase regulator activity
GO:0002376 0.0033 0.021 immune system process
GO:0007018 0.0034 0.021 microtubule-based movement
GO:0000041 0.0034 0.021 transition metal ion transport
GO:0006465 0.0037 0.022 signal peptide processing
GO:0005905 0.0065 0.022 coated pit
GO:0030132 0.0065 0.022 clathrin coat of coated pit
GO:0044237 0.0039 0.023 cellular metabolic process
GO:0008152 0.004 0.023 metabolic process
GO:0060589 0.0043 0.023 nucleoside-triphosphatase regulator activity
GO:0042558 0.0045 0.025 pteridine-containing compound metabolic process
GO:0006457 0.0045 0.025 protein folding
GO:0006106 0.0045 0.025 fumarate metabolic process
GO:0042398 0.0047 0.026 cellular modified amino acid biosynthetic process
GO:0016248 0.0056 0.028 channel inhibitor activity
GO:0008200 0.0056 0.028 ion channel inhibitor activity
GO:0016247 0.0063 0.03 channel regulator activity
GO:0039624 0.01 0.031 viral outer capsid
GO:0019030 0.011 0.031 icosahedral viral capsid
GO:0015629 0.011 0.031 actin cytoskeleton
GO:0030118 0.011 0.031 clathrin coat
GO:0031227 0.011 0.031 intrinsic to endoplasmic reticulum membrane
GO:0044763 0.0059 0.032 single-organism cellular process
GO:0005515 0.0067 0.032 protein binding
GO:0032318 0.0062 0.033 regulation of Ras GTPase activity
GO:0019028 0.012 0.033 viral capsid
GO:0009987 0.0066 0.035 cellular process
GO:0031012 0.015 0.04 extracellular matrix
GO:0071704 0.008 0.041 organic substance metabolic process
GO:0030030 0.0081 0.042 cell projection organization
GO:0031301 0.018 0.045 integral to organelle membrane
GO:0035251 0.01 0.045 UDP-glucosyltransferase activity
GO:0051701 0.0095 0.048 interaction with host
Cluster 4
term_id pvalue adjp term_name
GO:0015459 2.50E-10 1.30E-08 potassium channel regulator activity
GO:0019870 2.50E-10 1.30E-08 potassium channel inhibitor activity
GO:0016248 1.70E-09 4.20E-08 channel inhibitor activity
GO:0008200 1.70E-09 4.20E-08 ion channel inhibitor activity
GO:0016247 2.10E-09 4.20E-08 channel regulator activity
GO:0031423 5.00E-06 8.40E-05 hexon binding
GO:0004803 1.10E-05 0.00017 transposase activity
GO:0071944 1.40E-05 0.00065 cell periphery
GO:0005886 0.00024 0.0056 plasma membrane
GO:0030312 0.00052 0.0069 external encapsulating structure
GO:0044459 0.00059 0.0069 plasma membrane part
GO:0004672 8.00E-04 0.01 protein kinase activity
GO:0004713 0.0017 0.017 protein tyrosine kinase activity
GO:0008175 0.0017 0.017 tRNA methyltransferase activity
GO:0006298 8.40E-05 0.02 mismatch repair
GO:0019199 0.0028 0.026 transmembrane receptor protein kinase activity
GO:0042302 0.0037 0.031 structural constituent of cuticle
GO:0005905 0.0048 0.032 coated pit
GO:0030132 0.0048 0.032 clathrin coat of coated pit
GO:0030118 0.0069 0.032 clathrin coat
GO:0016011 0.007 0.032 dystroglycan complex
GO:0016012 0.007 0.032 sarcoglycan complex
GO:0034464 0.0072 0.032 BBSome
GO:0016010 0.0076 0.032 dystrophin-associated glycoprotein complex
GO:0038023 0.0048 0.037 signaling receptor activity
GO:0006281 0.00054 0.042 DNA repair
GO:0006811 0.00059 0.042 ion transport
GO:0006974 0.00081 0.042 response to DNA damage stimulus
GO:0043244 0.0016 0.042 regulation of protein complex disassembly
GO:0033554 0.0021 0.042 cellular response to stress
GO:0090407 0.0027 0.042 organophosphate biosynthetic process
GO:0072528 0.0028 0.042 pyrimidine-containing compound biosynthetic process
GO:0044093 0.0029 0.042 positive regulation of molecular function
GO:0043085 0.0029 0.042 positive regulation of catalytic activity
GO:0009233 0.0029 0.042 menaquinone metabolic process
GO:0009234 0.0029 0.042 menaquinone biosynthetic process
GO:0044765 0.0035 0.042 single-organism transport
GO:0051694 0.0038 0.042 pointed-end actin filament capping
GO:0042357 0.0044 0.042 thiamine diphosphate metabolic process
GO:0009229 0.0044 0.042 thiamine diphosphate biosynthetic process
GO:0051234 0.0045 0.042 establishment of localization
GO:0006810 0.0045 0.042 transport
GO:1901879 0.0049 0.042 regulation of protein depolymerization
GO:0031333 0.0049 0.042 negative regulation of protein complex assembly
GO:0030834 0.0049 0.042 regulation of actin filament depolymerization
GO:0032272 0.0049 0.042 negative regulation of protein polymerization
GO:0051494 0.0049 0.042 negative regulation of cytoskeleton organization
GO:1901880 0.0049 0.042 negative regulation of protein depolymerization
GO:0030835 0.0049 0.042 negative regulation of actin filament depolymerization
GO:0030837 0.0049 0.042 negative regulation of actin filament polymerization
GO:0051693 0.0049 0.042 actin filament capping
GO:0043242 0.0062 0.042 negative regulation of protein complex disassembly
GO:0045727 0.0062 0.042 positive regulation of translation
GO:0019645 0.0062 0.042 anaerobic electron transport chain
GO:0006835 0.0066 0.042 dicarboxylic acid transport
GO:0019637 0.007 0.042 organophosphate metabolic process
GO:0006796 0.007 0.042 phosphate-containing compound metabolic process
GO:0006793 0.0073 0.042 phosphorus metabolic process
GO:0072527 0.0073 0.042 pyrimidine-containing compound metabolic process
GO:0090066 0.0077 0.042 regulation of anatomical structure size
GO:0032535 0.0077 0.042 regulation of cellular component size
GO:0032970 0.0077 0.042 regulation of actin filament-based process
GO:0043254 0.0077 0.042 regulation of protein complex assembly
GO:0030832 0.0077 0.042 regulation of actin filament length
GO:0032271 0.0077 0.042 regulation of protein polymerization
GO:0032956 0.0077 0.042 regulation of actin cytoskeleton organization
GO:0030833 0.0077 0.042 regulation of actin filament polymerization
GO:0008064 0.0077 0.042 regulation of actin polymerization or depolymerization
GO:0004872 0.0062 0.045 receptor activity
GO:0004529 0.0072 0.045 exodeoxyribonuclease activity
GO:0016895 0.0072 0.045 “exodeoxyribonuclease activity, producing 5′-phosphomonoesters”
GO:0044087 0.0091 0.048 regulation of cellular component biogenesis
GO:0010639 0.0095 0.049 negative regulation of organelle organization
Cluster 5
term_id pvalue adjp term_name
GO:0015459 1.70E-13 8.70E-12 potassium channel regulator activity
GO:0019870 1.70E-13 8.70E-12 potassium channel inhibitor activity
GO:0016247 4.20E-13 1.40E-11 channel regulator activity
GO:0016248 2.00E-12 4.00E-11 channel inhibitor activity
GO:0008200 2.00E-12 4.00E-11 ion channel inhibitor activity
GO:0004336 0.00019 0.0032 galactosylceramidase activity
GO:0016301 0.00087 0.012 kinase activity
GO:0019866 0.00037 0.015 organelle inner membrane
GO:0016772 0.0016 0.018 “transferase activity, transferring phosphoruscontaining groups”
GO:0004114 0.0017 0.018 “3′,5′-cyclic-nucleotide phosphodiesterase activity”
GO:0004371 0.0021 0.021 glycerone kinase activity
GO:0005200 0.0033 0.03 structural constituent of cytoskeleton
GO:0031224 0.0044 0.035 intrinsic to membrane
GO:0016021 0.0048 0.035 integral to membrane
GO:0016011 0.005 0.035 dystroglycan complex
GO:0016012 0.005 0.035 sarcoglycan complex
GO:0016010 0.0053 0.035 dystrophin-associated glycoprotein complex
GO:0031967 0.0068 0.038 organelle envelope
GO:0042578 0.0055 0.046 phosphoric ester hydrolase activity
Cluster 6
term_id pvalue adjp term_name
GO:0016462 2.60E-07 8.20E-06 pyrophosphatase activity
GO:0016818 2.90E-07 8.20E-06 “hydrolase activity, acting on acid
anhydrides, in phosphorus-containing anhydrides”
GO:0016817 3.70E-07 8.20E-06 “hydrolase activity, acting on acid anhydrides”
GO:0016887 1.80E-06 2.60E-05 ATPase activity
GO:0017111 2.00E-06 2.60E-05 nucleoside-triphosphatase activity
GO:0006298 7.30E-07 8.10E-05 mismatch repair
GO:0015459 3.60E-05 0.00034 potassium channel regulator activity
GO:0019870 3.60E-05 0.00034 potassium channel inhibitor activity
GO:0004371 8.40E-05 0.00059 glycerone kinase activity
GO:0016248 8.80E-05 0.00059 channel inhibitor activity
GO:0008200 8.80E-05 0.00059 ion channel inhibitor activity
GO:0016247 9.60E-05 0.00059 channel regulator activity
GO:0018143 1.70E-05 7.00E-04 nucleic acid-protein covalent cross-linking
GO:0006811 2.00E-05 7.00E-04 ion transport
GO:0006281 2.50E-05 7.00E-04 DNA repair
GO:0006974 3.80E-05 0.00085 response to DNA damage stimulus
GO:0016011 0.00049 0.0042 dystroglycan complex
GO:0016012 0.00049 0.0042 sarcoglycan complex
GO:0016010 0.00053 0.0042 dystrophin-associated glycoprotein complex
GO:0045333 0.00031 0.0057 cellular respiration
GO:0010604 0.00056 0.0065 positive regulation of macromolecule metabolic process
GO:0033554 0.00061 0.0065 cellular response to stress
GO:0006259 0.00061 0.0065 DNA metabolic process
GO:0009893 0.00064 0.0065 positive regulation of metabolic process
GO:0031325 0.00064 0.0065 positive regulation of cellular metabolic process
GO:0008152 0.0013 0.012 metabolic process
GO:0015980 0.0014 0.012 energy derivation by oxidation of organic compounds
GO:0044237 0.0019 0.013 cellular metabolic process
GO:0006091 0.002 0.013 generation of precursor metabolites and energy
GO:0032270 0.002 0.013 positive regulation of cellular protein metabolic process
GO:0051247 0.002 0.013 positive regulation of protein metabolic process
GO:0042357 0.0021 0.013 thiamine diphosphate metabolic process
GO:0009229 0.0021 0.013 thiamine diphosphate biosynthetic process
GO:0055114 0.0023 0.013 oxidation-reduction process
GO:0009891 0.0026 0.013 positive regulation of biosynthetic process
GO:0010557 0.0026 0.013 positive regulation of macromolecule biosynthetic process
GO:0031328 0.0026 0.013 positive regulation of cellular biosynthetic process
GO:0016538 0.0023 0.013 cyclin-dependent protein serine/threonine kinase regulator activity
GO:0015399 0.0027 0.013 primary active transmembrane transporter activity
GO:0015405 0.0027 0.013 P-P-bond-hydrolysis-driven transmembrane transporter activity
GO:0046483 0.003 0.014 heterocycle metabolic process
GO:0048522 0.0042 0.019 positive regulation of cellular process
GO:0034641 0.0044 0.019 cellular nitrogen compound metabolic process
GO:0071944 0.0032 0.019 cell periphery
GO:0051716 0.0051 0.02 cellular response to stimulus
GO:0006950 0.0052 0.02 response to stress
GO:0044459 0.0054 0.022 plasma membrane part
GO:0005886 0.0055 0.022 plasma membrane
GO:1901360 0.006 0.023 organic cyclic compound metabolic process
GO:0006725 0.0063 0.023 cellular aromatic compound metabolic process
GO:0090304 0.0064 0.023 nucleic acid metabolic process
GO:0036211 0.0072 0.024 protein modification process
GO:0006464 0.0072 0.024 cellular protein modification process
GO:0072528 0.009 0.029 pyrimidine-containing compound biosynthetic process
GO:0044765 0.01 0.032 single-organism transport
GO:0048583 0.01 0.032 regulation of response to stimulus
GO:0048518 0.013 0.039 positive regulation of biological process
GO:0004803 0.011 0.04 transposase activity
GO:0016301 0.011 0.04 kinase activity
GO:0004861 0.011 0.04 cyclin-dependent protein serine/threonine kinase inhibitor activity
GO:0016668 0.012 0.04 “oxidoreductase activity, acting on a sulfur group of donors, NAD(P) as acceptor”
GO:0030291 0.012 0.04 protein serine/threonine kinase inhibitor activity
GO:0047134 0.012 0.04 protein-disulfide reductase activity
GO:0006139 0.014 0.041 nucleobase-containing compound metabolic process
GO:0051234 0.015 0.041 establishment of localization
GO:0006810 0.015 0.041 transport
GO:0072527 0.017 0.047 pyrimidine-containing compound metabolic process
GO:0039624 0.016 0.047 viral outer capsid
GO:0019028 0.017 0.047 viral capsid
GO:0019030 0.017 0.047 icosahedral viral capsid
Cluster 7
term_id pvalue adjp term_name
GO:0006298 1.20E-11 2.80E-09 mismatch repair
GO:0015459 1.40E-10 8.80E-09 potassium channel regulator activity
GO:0019870 1.40E-10 8.80E-09 potassium channel inhibitor activity
GO:0016247 3.80E-10 1.60E-08 channel regulator activity
GO:0016248 1.40E-09 3.40E-08 channel inhibitor activity
GO:0008200 1.40E-09 3.40E-08 ion channel inhibitor activity
GO:0004803 2.30E-09 4.70E-08 transposase activity
GO:0006974 7.30E-09 8.20E-07 response to DNA damage stimulus
GO:0006281 1.70E-08 1.30E-06 DNA repair
GO:0016011 1.40E-07 2.20E-06 dystroglycan complex
GO:0016012 1.40E-07 2.20E-06 sarcoglycan complex
GO:0016010 1.60E-07 2.20E-06 dystrophin-associated glycoprotein complex
GO:0044459 1.00E-06 8.90E-06 plasma membrane part
GO:0005886 1.10E-06 8.90E-06 plasma membrane
GO:0071944 5.20E-06 3.50E-05 cell periphery
GO:0033554 6.80E-07 3.80E-05 cellular response to stress
GO:0016887 1.50E-05 0.00026 ATPase activity
GO:0072528 9.20E-06 0.00041 pyrimidine-containing compound biosynthetic process
GO:0042357 2.20E-05 7.00E-04 thiamine diphosphate metabolic process
GO:0009229 2.20E-05 7.00E-04 thiamine diphosphate biosynthetic process
GO:0044237 4.60E-05 0.0011 cellular metabolic process
GO:0090407 4.90E-05 0.0011 organophosphate biosynthetic process
GO:0072527 5.10E-05 0.0011 pyrimidine-containing compound metabolic process
GO:0016818 6.80E-05 0.0011 “hydrolase activity, acting on acid anhydrides, in phosphorus-containing anhydrides”
GO:0016817 8.80E-05 0.0012 “hydrolase activity, acting on acid anhydrides”
GO:0006950 6.60E-05 0.0013 response to stress
GO:0017111 0.00012 0.0015 nucleoside-triphosphatase activity
GO:0051716 8.90E-05 0.0016 cellular response to stimulus
GO:0016462 0.00017 0.0019 pyrophosphatase activity
GO:0046483 0.00019 0.0032 heterocycle metabolic process
GO:0034641 0.00021 0.0032 cellular nitrogen compound metabolic process
GO:0008152 0.00022 0.0032 metabolic process
GO:0006811 0.00026 0.0036 ion transport
GO:0006725 0.00041 0.0054 cellular aromatic compound metabolic process
GO:0045333 0.00051 0.0063 cellular respiration
GO:0006357 0.00057 0.0066 regulation of transcription from RNA polymerase II promoter
GO:0031225 0.0012 0.007 anchored to membrane
GO:1901360 7.00E-04 0.0074 organic cyclic compound metabolic process
GO:0006790 7.00E-04 0.0074 sulfur compound metabolic process
GO:0044272 0.00094 0.0095 sulfur compound biosynthetic process
GO:0015980 0.001 0.01 energy derivation by oxidation of organic compounds
GO:0019637 0.0012 0.011 organophosphate metabolic process
GO:0006796 0.0012 0.011 phosphate-containing compound metabolic process
GO:0006259 0.0014 0.011 DNA metabolic process
GO:0010604 0.0014 0.011 positive regulation of macromolecule metabolic process
GO:0009893 0.0016 0.012 positive regulation of metabolic process
GO:0019438 0.0016 0.012 aromatic compound biosynthetic process
GO:0031325 0.0016 0.012 positive regulation of cellular metabolic process
GO:0006091 0.0017 0.012 generation of precursor metabolites and energy
GO:0006732 0.0018 0.012 coenzyme metabolic process
GO:0032784 0.0018 0.012 “regulation of DNA-dependent transcription, elongation”
GO:0009108 0.002 0.013 coenzyme biosynthetic process
GO:0055114 0.0021 0.013 oxidation-reduction process
GO:0018130 0.0023 0.014 heterocycle biosynthetic process
GO:0006355 0.0031 0.019 “regulation of transcription, DNA-dependent”
GO:0006793 0.0034 0.02 phosphorus metabolic process
GO:0044271 0.0035 0.02 cellular nitrogen compound biosynthetic process
GO:0031423 0.002 0.021 hexon binding
GO:0051188 0.0039 0.022 cofactor biosynthetic process
GO:0051186 0.0043 0.023 cofactor metabolic process
GO:0051252 0.0044 0.023 regulation of RNA metabolic process
GO:2001141 0.0044 0.023 regulation of RNA biosynthetic process
GO:0032270 0.0052 0.026 positive regulation of cellular protein metabolic process
GO:0051247 0.0052 0.026 positive regulation of protein metabolic process
GO:0090304 0.0055 0.027 nucleic acid metabolic process
GO:0018143 0.0056 0.027 nucleic acid-protein covalent cross-linking
GO:1901362 0.0059 0.027 organic cyclic compound biosynthetic process
GO:0004861 0.003 0.029 cyclin-dependent protein serine/threonine kinase inhibitor activity
GO:0016538 0.0036 0.03 cyclin-dependent protein serine/threonine kinase regulator activity
GO:0030291 0.0036 0.03 protein serine/threonine kinase inhibitor activity
GO:0006139 0.0068 0.031 nucleobase-containing compound metabolic process
GO:0071822 0.0072 0.031 protein complex subunit organization
GO:0000226 0.0075 0.031 microtubule cytoskeleton organization
GO:0032786 0.0075 0.031 “positive regulation of DNA-dependent transcription, elongation”
GO:0034243 0.0075 0.031 regulation of transcription elongation from RNA polymerase II promoter
GO:0032968 0.0075 0.031 positive regulation of transcription elongation from RNA polymerase II promoter
GO:0006221 0.0097 0.039 pyrimidine nucleotide biosynthetic process
GO:0060255 0.01 0.041 regulation of macromolecule metabolic process
GO:0010468 0.011 0.041 regulation of gene expression
GO:0008175 0.0053 0.041 tRNA methyltransferase activity
GO:0070035 0.006 0.041 purine NTP-dependent helicase activity
GO:0008026 0.006 0.041 ATP-dependent helicase activity
GO:0004371 0.007 0.044 glycerone kinase activity
GO:0042623 0.0071 0.044 “ATPase activity, coupled”
GO:0006220 0.012 0.048 pyrimidine nucleotide metabolic proces
~~~

